# Drastic improvement in MHC class I epitope prediction through training data augmentation, generative adversarial network modeling of selfness, and consideration of homology to class II epitopes

**DOI:** 10.64898/2026.11.16.766890

**Authors:** Sejin Oh, Seong-Eui Hong, Hae-Suk Kim, Soonmyung Paik

## Abstract

Accurate identification of immunogenic neoantigens is critical for personalized cancer vaccines but remains hindered by the high false-positive rates of current predictors. Here, we present DeepOmicsNeo, a modular deep learning framework that harmonizes three biological determinants of immunogenicity. The first module, DeepOmicsNeo-Binding, overcomes data scarcity in non-9mer peptide-Class I Major Histocompatibility Complex (MHC) binding prediction via a novel augmentation strategy that synthesizes pseudo-binders and non-binders by editing non-anchor residues, outperforming its competitors. The second module, DeepOmicsNeo-Selfness, addresses the paucity of negative training data for immunogenicity prediction by utilizing a Generative Adversarial Network trained on around 60 million human self-peptides. This unsupervised approach quantifies selfness to detect immunogenic peptides and the resulting score correlated with IFN-γ secretion by T-cells. The third module, DeepOmicsNeo-H2, exploits the central-region homology between known HLA Class-I and Class-II restricted epitopes. The integrated model, DeepOmicsNeo-Immunogenicity, outperforms state-of-the-art predictors on the TESLA, HiTIDE, and TiTANIAN benchmark datasets. Top20 precision, the most important benchmark for clinical utility in personalized cancer vaccine design, was drastically improved over other predictors. Together, these findings indicate that jointly modeling the three complementary features captures biologically meaningful determinants of immunogenicity, supporting DeepOmicsNeo as a robust computational framework for neoantigen prioritization in personalized cancer immunotherapy.

## Introduction

For personalized cancer vaccine design, selection of 20 to 30 potentially immunogenic neoepitopes derived from somatic mutations using a computational prediction algorithm is required^1,2^. However, currently available prediction algorithms suffer from low top20 precision with a high false positivity, i.e. many of the top 20 selected candidates are not immunogenic^3,4^.

Effective priming of a CD8+ T-cell-mediated immune response by a potentially immunogenic mutant peptide requires the fulfillment of several critical conditions^5^: 1) MHC Class I presentation: The peptide must bind to a Class I Major Histocompatibility Complex (MHC) protein and be presented as peptide-MHC complex (pMHC) on the surface of antigen-presenting cells, such as dendritic cells. 2) Naive T-cell availability: Naive T-cells possessing cognate T-cell receptors (TCRs) recognizing the pMHC must be present within the secondary lymphoid organs draining the tumor tissue. 3) CD4+ T Cell help: Antigen-presenting cells require assistance from CD4+ helper T cells (forming the APC-CD4-CD8 immune triad) to facilitate the full differentiation of naive T cells into effector T cells. We hypothesized that a model integrating these concepts may greatly improve the top20 precision when designing personalized cancer vaccines.

However, significant barriers currently impede the development of an integrated model with a high precision. 1) MHC binding affinity data limitations: While predictions of peptide-MHC binding affinity are generally robust, accuracy for non-9mer peptides remains suboptimal due to the limited amount of training data available^6^. 2) TCR Repertoire Complexity and lack of training data: Cataloging an individual’s complete TCR repertoire is currently not feasible^7^.

Furthermore, developing a generalizable predictive model for pMHC-TCR binding is hindered by sparse data availability and the inherent cross-reactivity and degeneracy of TCRs^8^. 3) Absence of prediction models for CD4 Help: There are currently no predictive frameworks available that can infer CD4+ T cell help based solely on a Class I peptide-MHC sequence, although there are models that predict MHC class II epitopes.

We developed solutions addressing each of the above challenges and integrated them into a single predictive model DeepOmicsNeo for immunogenicity. 1) The first module, DeepOmicsNeo-Binding (DON-B), overcomes data scarcity in non-9mer peptide-MHC binding prediction via a novel augmentation strategy that synthesizes pseudo-binders by editing non-anchor residues, outperforming its competitors. 2) The second module, DeepOmicsNeo-Selfness (DON-S), is based on the concept that our adaptive immune system is never trained with foreign antigens and the TCR repertoire (armed with cross reactivity and degeneracy) that has a duty to recognize foreign antigens remains after self-reactive TCRs are eliminated in the thymus. Thus a Generative Adversarial Network trained only with normal human peptidome may serve as a surrogate for our self-reactive eliminated TCR repertoire and a peptide with low selfness score will likely have cognate TCRs available. This unsupervised approach quantifies selfness to detect immunogenic peptides and the resulting score correlated with functional avidity and immunogenicity. Use of a generative model addresses the problem of paucity of training data for immunogenicity. 3) The third module, DeepOmicsNeo-H2 (DON-H2), exploits the central-region homology between known HLA Class-I and Class-II restricted epitopes.

The integrated model, DeepOmicsNeo-Immunogenicity (DON-IM), integrating the above three components at the network structure level and trained ground up, outperforms state-of-the-art predictors on the TESLA, HiTIDE, and TiTANIAN benchmark datasets. We demonstrate that Top20 precision, the most important benchmark for clinical utility in personalized cancer vaccine design, was drastically improved over other predictors.

## Method

### Training and test datasets

Table 1 lists the datasets used for the training and testing of DeepOmicsNeo models. Briefly, we used a peptide-MHC-I (pMHC-I) binding dataset from the Immune Epitope Database (IEDB)^9^ in 2024, five public mono-allele immunopeptidome datasets^10–14^, to train DeepOmicsNeo-Binding. For DeepOmicsNeo-Selfness, we used an 8-12mer sliced human reference peptidome dataset sourced from Ensembl^15^. For DeepOmicsNeo-Homology-with-HLA-II-epitope, we used the T-cell response dataset of IEDB (IEDB-T). For DeepOmicsNeo-Immunogenicity, we used IEDB-T and a preprocessed neoantigen dataset^16^.

**Table 1.** Used datasets in this study.

| Category | Name | Used model | Reference |
|---|---|---|---|
| Training | IEDB (pMHC-I binding) | DON-B | <sup>9</sup> |
|  | Mono-alleles | DON-B | <sup>11–14</sup> |
|  | Human reference peptidome | DON-S | <sup>15</sup> |
|  | IEDB (T-cell response) | DON-IM, DON-H2 | <sup>9</sup> |
|  | NCI | DON-IM | <sup>16</sup> |
| Test | Binding benchmark with multi-allelic immunopeptidome datasets | DON-B<br>DON-IM<br>NetMHCpan (v4.1b)<br>MHCflurry (v2.0)<br>MHCnuggets (v1.0)<br>MixMHCpred (v3.0)<br>PRIME (v2.1)<br>BigMHC (v1.0) | <sup>12,14,17–19</sup> |
|  | TESLA | All models | <sup>3</sup> |
|  | HiTIDE | DON-B<br>DON-IM<br>NetMHCpan (v4.1b)<br>MHCflurry (v2.0)<br>MHCnuggets (v1.0)<br>MixMHCpred (v3.0)<br>PRIME (v2.1)<br>BigMHC (v1.0) | <sup>16</sup> |
|  | TiTANIAN-neoantigen | DON-B<br>DON-IM<br>NetMHCpan (v4.1b)<br>MHCflurry (v2.0)<br>MHCnuggets (v1.0)<br>MixMHCpred (v3.0)<br>PRIME (v2.1)<br>BigMHC (v1.0) | <sup>20</sup> |
|  | HLA-agnostic immune response | DON-S | <sup>21</sup> |
|  | T-cell avidity | DON-S | <sup>22</sup> |

To ensure a strict separation between training and testing, all samples appearing in the test datasets were excluded from the training datasets.

Test datasets are composed of five multi-allelic immunopeptidome datasets, TESLA, HiTIDE, TiTANIAN, and HLA-agnostic dataset. Details of the datasets are described in the following sections.

### pMHC-I training dataset and DeepOmicsNeo-Binding model training

In this study, the training and test datasets for peptide-MHC class I (pMHC-I) binding were sourced from the Immune Epitope Database (IEDB)^9^ and five mono-allelic immunopeptidome datasets (MAIDs)^10–14^.

We specifically curated experiments involving 8 to 12mer peptides, resulting in a training dataset of 856,736 data points (realSet). For pMHC-Is without affinity values, we set surrogate-affinity (sAff) according to a method proposed by O’Donnell et al., with some modification^23^. We set 100, 500, 1,000, 5,000 and 10,000 nM as sAffs for ‘positive-high’, ‘positive or MAID’, ‘positive-intermediate’, ‘positive-low’, and ‘negative’ labels. For these pMHC-Is with sAffs, we applied a conditional-mean-squared-error (cMSE) loss function. During training, these pMHC-Is contribute to the MSE loss only when their predicted affinities (pAff) are greater (for positive binders, Direction (D) = -1) or less (for negative binders, D = +1) than their specified sAffs (Equation 1 and 2).

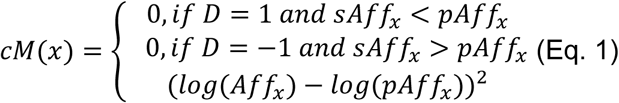

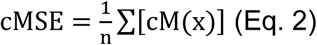

Although the cMSE loss function works, we noticed that this may cause insufficient training of the model likely due to artificial gradient vanishing. To mitigate this issue, we implemented weighted binary cross-entropy (wBCE) loss. We transformed the pAff of each pMHC-I sample x into a binding probability using Eq. 3. Next, the probabilities were converted into cross-entropy values using Eq. 4, where n is the batch size, and y is the binary label (1 for positive and 0 for negative binders). For probabilities from positive labels, we normalized the cross-entropy values by their sAffs to incorporate prior knowledge about qualitative characteristics. In contrast, cross-entropy values from negative labels were not normalized.

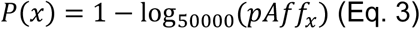

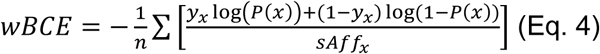

DeepOmicsNeo-Binding is trained with the sum of the two loss functions (Eq. 5).

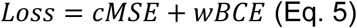

### Sequence similarity between pMHC-I binding peptides across peptide lengths

To evaluate whether pMHC-I binding peptides of different lengths show sequence similarity to the 9mer binder repertoire of the same MHC-I allele, we defined a dissimilarity metric, dissimilarity-to-self (Dis2self)^24^. For 8mer and 10mer peptides, each candidate insertion/deletion position (IDP) was evaluated by modifying the peptide into a 9-position alignment - inserting a blank at the candidate position for 8mers, or removing the amino acid at the candidate position for 10mers. For example, an 8mer AACCDDEE with a blank inserted at position 1 becomes -AACCDDEE, and a 10mer AACCDDEEFF with the amino acid at position 1 deleted becomes ACCDDEEFF. For a given IDP-modified peptide, Dis2self against a reference 9mer binder was calculated as the proportion of mismatched amino acids among all non-blank aligned positions; positions aligned to a blank were excluded from the comparison and did not contribute to the mismatch count.

For each IDP-modified peptide, the minimum Dis2self across all 9mer binders of the same MHC-I allele was retained, and among all candidate IDPs, the one yielding the lowest minimum Dis2self was selected as the best alignment for that peptide.

For 9mer peptides, since candidate and reference peptides are of equal length, no blank insertion was required. Dis2self for each 9mer binder was calculated as the direct position-wise calculation against every other 9mer binder of the same allele (excluding the peptide itself). The identical procedure was applied to random peptide sets generated in numbers equal to the real 8mer, 9mer, and 10mer binder sets, to obtain the background Dis2self distribution. Statistical significance between the Dis2self distributions of binder-derived peptides and length-matched random peptides was assessed using a two-sided Mann-Whitney U test. For visualization, Dis2self values were converted to a similarity score as 100 – Dis2self, such that higher values indicate greater sequence similarity to the allele’s 9mer binder repertoire.

For 8mer, 9mer, and 10mer peptides, the positional distribution of edits (insertions/deletions, for 8mer/10mer) or mismatches (for 9mer, derived from the Dis2self alignment) yielding the maximum alignment score against the most similar 9mer was examined. Edit/mismatch frequency at each peptide position was calculated across all binder-allele pairs and compared against the corresponding frequency in random peptide sets.

### Data augmentation for pMHC-I binding dataset

We generated a negative binding dataset (negSet) 100 times larger than the realSet by incorporating random peptides. To construct an augmented positive dataset (augSet) for training of a K-mer affinity prediction model, pMHC-I with strong affinities (< 1,000nM) of three lengths were generated from the realSet: K−1 mer peptides were converted by inserting a single random amino acid, K-mer peptides by substituting a single random amino acid, and K+1 mer peptides by deleting a single random amino acid, all within the general anchor positions (P3 to Pω−1). For example, to train a 9mer affinity prediction model, the augSet was constructed by converting peptides of neighboring lengths into 9mers: 8mer peptides were extended to 9mers by single-amino-acid insertion, 9mer peptides were retained as 9mers with single-amino-acid substitution, and 10mer peptides were trimmed to 9mers by single-amino-acid deletion.

The affinity labels for peptides in the augSet and negSet were revised using the editor model trained with the realSet and negSet (Figure 2A).

### DeepOmicsNeo-Binding model

DeepOmicsNeo-Binding is composed of aligners and affinity-predictors. The aligners return bindable (bP9) and non-bindable 9-mer profiles (nP9) for non-9mer peptides for a given MHC-I allele by calculating the probability of each insertion/deletion position (IDP) in the peptides. The affinity predictors calculate affinities of pMHC-I from bP9, nP9, full peptide sequence, and pseudo-34mer MHC-I sequences. Peptide sequences are encoded by BLOSUM62 values. The activation function and optimizer of the models are the Gaussian error linear unit (GELU) and Nesterov-accelerated adaptive moment estimation (NAdam) optimizer. The model consists of two fully connected residual blocks with stochastic depth, where the survival probability decays linearly with depth^25^, from 0.9 in the first block to 0.8 in the second (Supplementary Figure 1D). All models were trained by 10-fold cross-validation with a batch size of 1024. All models were implemented in PyTorch (version 2.6.0). Details are described in the following sections.

**Figure S1.**
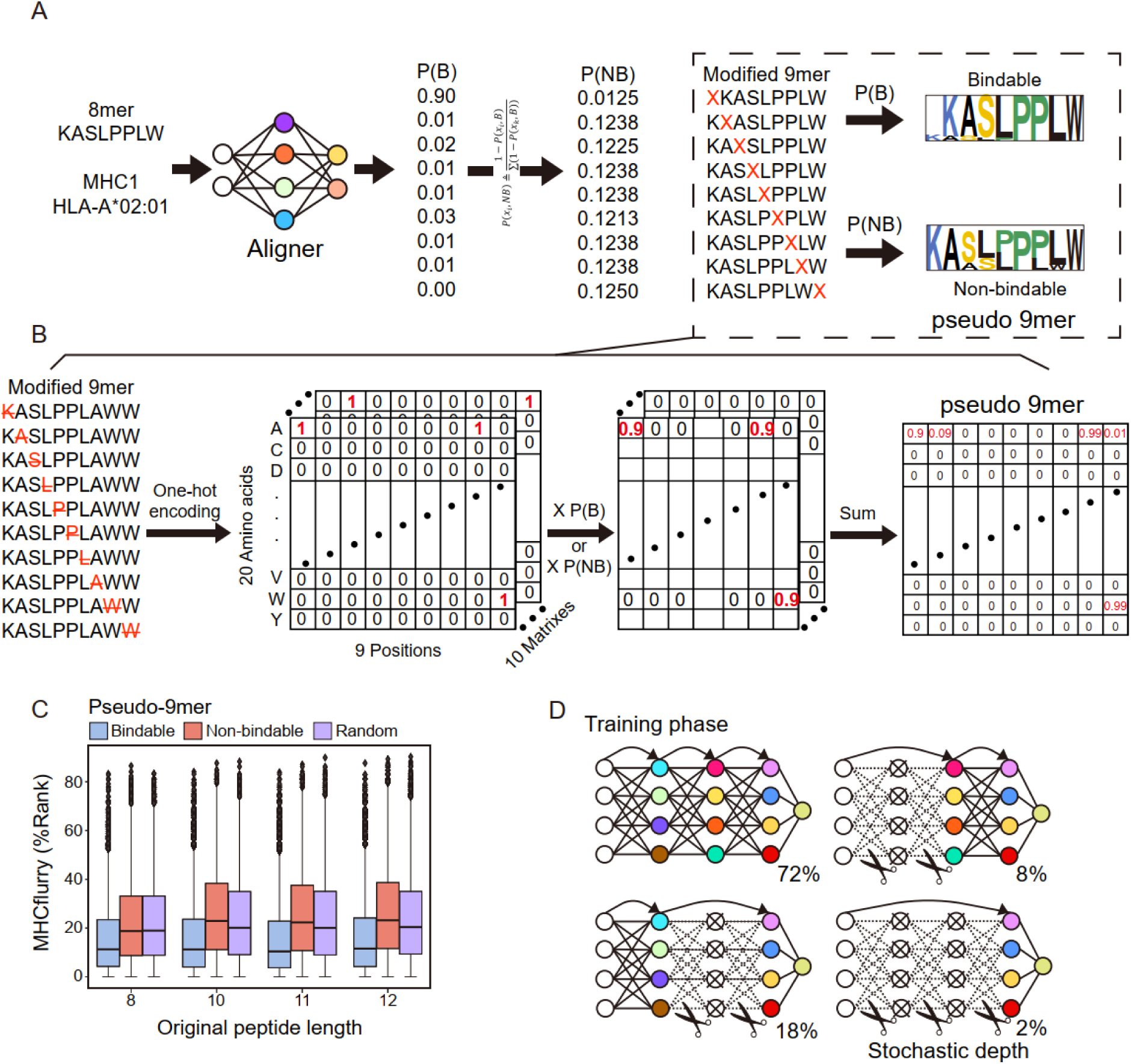
Details of pseudo-9mer generation and DeepOmicsNeo-Binding training. A and B. Overall workflow of how aligners generate pseudo-9mers. For a given non-9mer peptide sequence and an MHC-I allele, aligners score positions on the peptide. Higher scores indicate modification of the positions will make the peptide bindable to the MHC-I allele. The scores (probability scale) are used to generate bindable and non-bindable pseudo 9mer profiles of the peptide. See Methods for details. C. Comparison of predicted affinities between bindable and non-bindable pseudo-9mers. The Y-axis represents the percentile rank (%Rank) of pMHC-I affinity predicted by MHCflurry, where lower values indicate stronger binding affinity. The legends are the same as Figure 1B. Exact p-values are detailed in Supplementary Table 1. D. Layer statuses of DeepOmicsNeo-Binding during its training phase. Circles with “X” and dotted lines represent dropped layers and disconnections to subsequent layers. The proportions of layer statuses are shown as percentages.

### Data preparation and training for aligner

We extracted 9mer pMHC-Is with affinities of 500nM or less from realSet. Next, we clustered MHC-I alleles based on their pseudo-34mer sequences and selected 154 MHC-I clusters containing at least 50 unique peptides each to construct 20-by-9 Position-propensity-matrixes (PPMs). We generated 1 million random peptides for each K-mer and computed the insertion position (8mer) or deletion position (from 10 to 12mer) where the alignment score of the modified peptide is maximized. Let MP(x) be the set of all possible modified peptides from non-9mer peptide x (Eq. 6). IDP(x) is the set of insertion/deletion positions for the given peptide x.

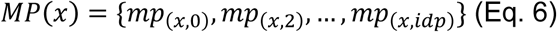

The alignment score of mp for a given MHC-I is calculated by Hadamard product of one-hot encoded mp and PPM (Eq. 7).

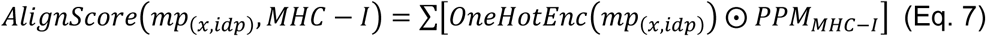

Finally, we selected the IDP with the highest alignment score as the aligner’s answer (Eq. 8).

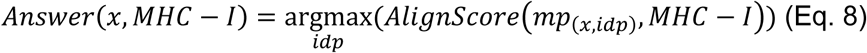

As the length of the peptide increases, the number of possible IDP combinations grows significantly. Therefore, when predicting IDPs, we limited our answers to continuous amino acid sequences. For example, in a 12mer peptide, deleting any 3 amino acids results in 220 possible combinations. However, restricting the deletion to consecutive 3-amino-acid segments reduces the number of possible combinations to 10, significantly decreasing the computational workload. Our aligners receive BLOSUM62 encoded non-9mer peptide sequence and pseudo-34mer sequence of MHC-I. Aligner models have varying numbers of neurons and layers depending on the input peptide length. The output layer uses a softmax function, allowing the models to predict answers based on probability. Training continued until no further decrease in loss was observed on the validation set. We selected the model from the epoch where the lowest cross-entropy loss was achieved as our aligner model.

Transforming non-9mer to 9mer profiles

For a given pMHC-I, aligners calculate probabilities of IDPs. As the aligners are trained to locate IDPs with maximum alignment scores for PSSMs, the probabilities were used to obtain a bindable 9mer profile. In the case of non-bindable 9mer profile, we calculated non-binding probability from the probability by Eq. 9.

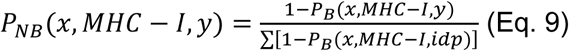

Finally, the bindable and non-bindable 9mer profiles for non-9mer peptides are calculated by Eq. 10.

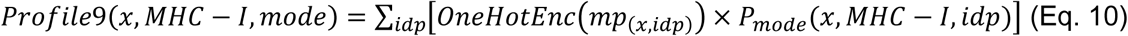

where mode indicates binding (B) or non-binding (NB) (Supplementary Figure 1A and 1B). The bP9 and nP9 profiles are each multiplied by the BLOSUM62 matrix (20 × 20) to yield pseudo-9mers (20 × 9), which serve as inputs to the affinity predictors.

**Figure 1.**
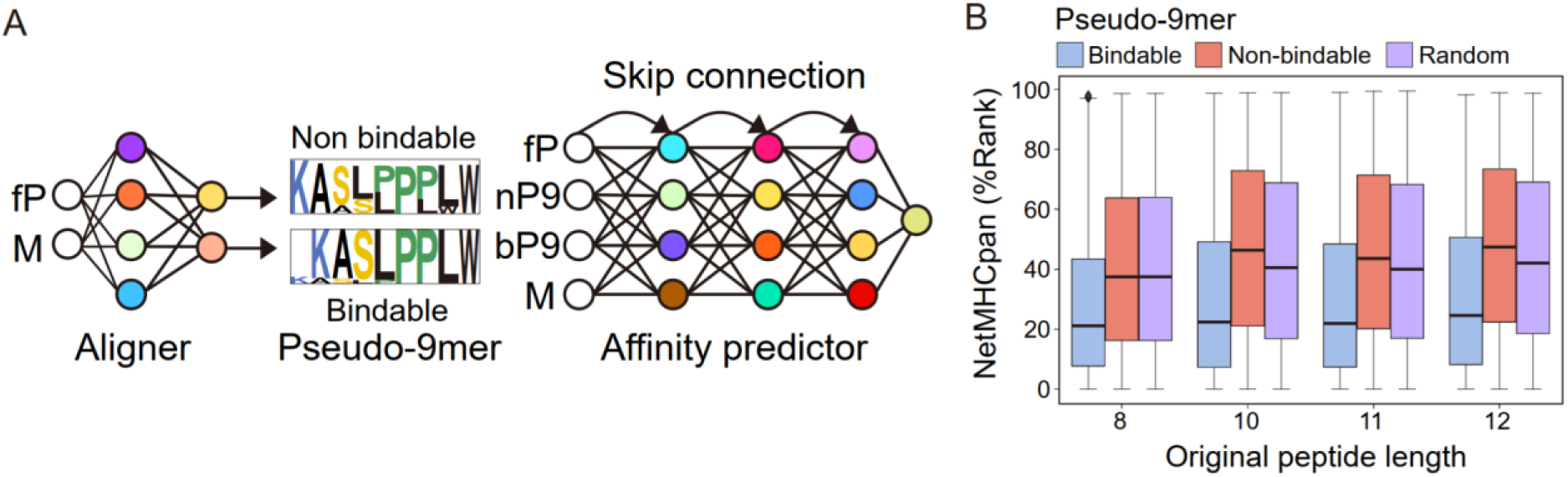
Structure of DeepOmicsNeo-Binding for non-9mer affinity prediction. A. Structure of non-9mer DeepOmicsNeo-Binding (DON-B). The aligner generates bindable (bP9) and non-bindable pseudo-9mers (nP9) from the full peptide sequence (fP) and MHC-I sequence (M). The affinity predictor predicts the pMHC-I affinity from fP, bP9, nP9, and M. See Methods for further details. B. Comparison of predicted affinities between bindable and non-bindable pseudo-9mers. The Y-axis represents the percentile rank (%Rank) of pMHC-I predicted by NetMHCpan, where lower values indicate stronger binding affinity. The X-axis represents the original peptide length. Blue, red, and purple boxplots indicate bindable, non-bindable, and random pseudo-9mer pMHC-Is, respectively. The box-and-whisker plots depict the medians (center line), first quartiles (lower bound), third quartiles (upper bound), and ±1.5× interquartile ranges (whiskers). Exact p-values are detailed in Supplementary Table 1.

**Figure 2.**
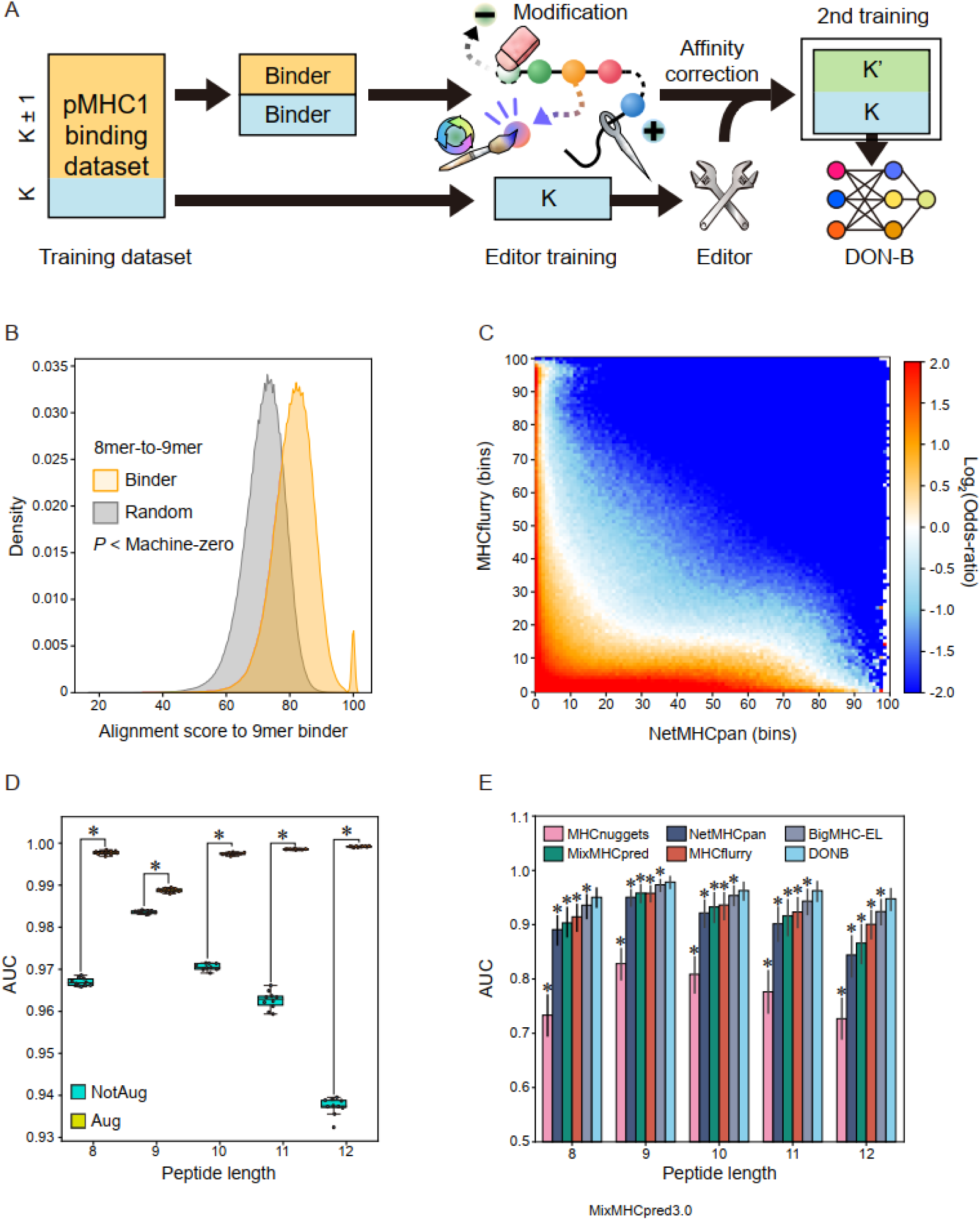
pMHC-I-optimized data augmentation. A. Workflow for training DeepOmicsNeo-Binding. To train a K-mer specific affinity predictor, the pMHC-I dataset of K and non-K-mers (differing from K-mers by +1 or -1 amino acid colored yellow) was modified to achieve K-mer length through insertion, substitution or deletion. The affinities of the modified peptides (K’) are estimated by the Editor models which were trained on K-mer and random-negative sets. Subsequently, DeepOmicsNeo-Binding (DON-B) was trained with both the corrected K’ (green) and K-mer (blue) datasets (See Methods for further details). B. Comparison of alignment scores between random- and binding-8mer peptides to binding-9mer peptides. Gray and yellow density plots illustrate the alignment score distributions for random-8mer sequences and pMHC-I binding 8mer sequences, respectively. C. Distribution of predicted binding affinities of augmented pMHC-I pairs derived from the binders. The X- and Y-axes represent the predicted binding affinity by NetMHCpan and MHCflurry, respectively, with values corresponding to percentiles (1–100). The color-scale bar indicates the Log_2_(odds-ratio) of the occurrence probability between the positive-binder-augmented dataset and the random-pMHC-I dataset. The binding affinities predicted by each tool were divided into 100 bins (percentiles). The Log_2_(odds-ratio) was calculated per bin by comparing the occurrence probability in the positive-binder-augmented dataset against the probability in the random-pMHC-I dataset. Positive values (red/orange) indicate that the augmented binder-derived data is significantly enriched in that specific affinity bin compared to the random pMHC-I pairs.ssss D. Performance comparison between models with and without data augmentation. The X-axis represents peptide sequence length, and the Y-axis represents the area under the curve (AUC) from 10-fold cross-validation. Models are color-coded: blue (trained without data augmentation) and yellow (trained with data augmentation). Asterisks indicate statistical significance (P < 0.05). Exact p-values are provided in Supplementary Table 2. E. Comparison of six pMHC-I binding prediction algorithms in multi-allelic benchmark sets. The X-axis represents peptide length, and the Y-axis represents the AUC score. Models are differentiated by color. Asterisks above barplots indicate that DON-B’s AUC values are statistically higher than those of the compared models. P-values were obtained by the Wilcoxon signed-rank test. Exact p-values are provided in Supplementary Table 2.

### Assessment of pseudo-9mer affinity

Random peptides of 8, 10, 11, and 12 residues were generated by sampling the 20 standard amino acids, with 1,000 peptides per length. The random seed was fixed at 20261103. For each HLA class I locus (HLA-A, -B, and -C), the five most frequent alleles in Koreans were selected based on allele frequencies reported by Park et al.^26^, giving 15 alleles in total. Each non-9mer peptide was converted into 9mer profiles using the aligner. For 8mers, alanine was inserted at the position with the maximum P(B) and, separately, at the position with the minimum P(B), yielding a bindable 9mer and a non-bindable 9mer. For peptides of 10 residues or longer, consecutive residues were deleted at the positions with the maximum and minimum P(B). For random 9mers, the modification site was chosen at random from positions other than the maximum and minimum P(B) indices, and the same insertion or deletion procedure was applied. Binding affinities of the resulting 9mers were predicted with NetMHCpan (v4.1b) and MHCflurry (v2.0) and compared as percentile ranks. Comparisons were performed for each K-mer length and for all peptides combined, for three pairs of groups: bindable vs. non-bindable, bindable vs. random, and non-bindable vs. random. Two-sided Mann–Whitney U tests were run using the scipy library (version 1.10.1).

### Generation of human self-peptidome

We collected a total of 121,766 human protein sequences from Ensembl (GRCh38 version 109)^15^. These protein sequences were broken down into peptides, each consisting of 8 to 12 amino acids. We excluded peptides containing ‘U’ (selenocysteine) or more than one ‘X’ (any amino acid). Subsequently, we replaced ‘X’ with 20 standard amino acids, generating around 60 million unique peptide sequences (11,343,860 for 8mer, 11,476,520 for 9mer, 11,560,513 for 10mer, 11,625,230 for 11mer, and 11,677,389 for 12mer).

### DeepOmicsNeo-Selfness model

DeepOmicsNeo-Selfness consists of five sub-models, each trained on different peptide lengths, respectively. They receive BLOSUM62 encoded peptide sequence (20 x K dimension, 20*K) and return selfness score. The generator consists of three layers employing two leaky-ReLU activations and batch normalization. The discriminator also comprises a similar structure to the generator. Our models were trained using the Wasserstein gradient penalty loss function, NAdam optimizer, 1,000 epochs, and a batch size of 1,024. We evaluated the model’s performance by assessing its ability to distinguish between human self-peptidome and a random peptidome using the AUC value. The random peptidome consisted of randomly generated peptides and was 10% of the size of the self-peptidome. We selected the discriminator from the epoch where the highest AUC was achieved as our DeepOmicsNeo-Selfness.

### Assessment of distinction ability between self- and nonself-peptidome

The reference protein sequences of model organisms were retrieved from the UniProt database^27^ using the following keywords: virus, chimpanzee, bacteria, and mouse. We selected pathogenic organisms listed in Annex 3 of the Pathogens Prioritization 2024 report by the World Health Organization. The protein sequences were processed similarly to human self-peptidome processing; however, sequences containing ‘OX=9606’ were removed, as these sequences are likely the exploited human proteins under infection rather than the organisms’ own proteins. Next, we calculated the selfness scores for each organism by averaging their peptide scores. The latent vectors of peptides were extracted from the last hidden layer of the DeepOmicsNeo-Selfness models. We used the averaged latent vector values as the representative peptidome vector for each organism. The distance between the human peptidome and the peptidome of each organism was calculated using the averaged latent vector.

### Assessment of correlation between DeepOmicsNeo-Selfness score and T-cell functional avidity or immunogenicity

T-cell functional avidity (EC50) dataset was obtained from Schmidt et al.^22^. The average of logarithms of EC50 values from ELISPOT assay were used to conduct Spearman’s correlation two-sided test using spearmanr function in scipy library (version 1.10.1). The HLA agnostic immunogenicity assay result was downloaded from Miller et al.^21^. We selected ELISPOT assay results (for IFN-γ and IL-5) obtained from missense variants and sliced them into short peptides, 8 to 12 amino acids in length, containing variant sites to make them compatible with our DeepOmicsNeo-Selfness. We set wildtype peptides as non-immunogenic peptides.

### DeepOmicsNeo-Homology-with-HLA-II-epitope model

8-12mer HLA-I epitopes were aligned against immunogenic HLA-II epitopes using local alignment in BLAST+ (parameters: -num_alignments 1, -num_threads 10, -evalue 1e10, - threshold 1, -word_size 2, -comp_based_stats 0, -seg no) and global alignment in ggsearch36 (parameters: -m 8CB, -s BL62, -E 1e10, -T 10, -b 1), with both tools configured to maximize alignment sensitivity. For each HLA-I epitope, the best-matching HLA-II epitopes was independently identified from the local and global alignment results. For HLA-I epitopes that could not be aligned to any HLA-II epitope by either tool, a supplementary alignment step was performed using Biopython (version 1.81): each unaligned HLA-I epitope was aligned against all candidate HLA-II epitopes, and the pair yielding the highest alignment score was selected as the matched pair. HLA-II epitopes were defined as immunogenic based on positive-assay labels in the source dataset.

From the resulting alignments, BLOSUM62 amino acid substitution scores between each HLA-I peptide (length A) and HLA-II peptide (length B) were used to construct an A-by-B substitution score matrix for each pair, with matrix dimensions varying according to the native lengths of the aligned peptides; variable-length matrices were grouped into batches via length-based bucketing during training. These matrices were used as input to a Squeeze- and-Excitation Network (SE-Net)^28^ comprising seven convolutional blocks (3×3 kernels; channel progression of 1, 32, 64, 128, 128, 128, 128, 128 across the network) with a squeeze-and-excitation reduction ratio of 4, followed by global average pooling. GELU activation functions were used throughout the network, which output a single logit representing the likelihood of immunogenicity; a sigmoid function was applied to this logit to obtain a probability. The model was trained using binary cross-entropy with logits loss (BCEWithLogitsLoss) and optimized with NAdam, using a batch size of 1024. Logits could be computed from substitution matrices derived from either the local- or global-alignment-based pairs; however, logits derived from local-alignment (BLAST+) pairs were used as the primary basis for this study, as the higher sensitivity of local alignment yielded a larger number of matched training pairs. Training was conducted on the IEDB-T dataset using 10-fold random-split cross-validation, and the model achieving the highest area under the curve (AUC) on the validation set in each fold was selected.

### Assessment of similarity between epitopes of HLA-I and HLA-II

To assess overall sequence similarity, we used HLA-II epitopes equal or longer than 15 amino acids as target sequences. HLA-I epitopes and length-matched peptides randomly sampled from the human wildtype peptidome were aligned against these HLA-II epitopes using BLAST+, ggsearch, and Biopython, as described above. For each query HLA-I epitope, the highest-scoring HLA-II epitope was selected independently for local and global alignments.

For positional Δscore analysis, we restricted the analysis to immunogenic HLA-I 9-mer epitopes and HLA-II 15-mer epitopes. For each immunogenic HLA-I 9-mer epitope, the highest-scoring HLA-II 15-mer epitope was independently identified from the immunogenic and non-immunogenic HLA-II epitope sets using local and global alignment. Thus, each HLA-I epitope could be associated with up to four HLA-II epitopes: one immunogenic and one non-immunogenic HLA-II epitope for each alignment method. For each selected HLA-I and HLA-II epitope pair, local and global alignments were performed using Biopython with an affine gap-opening penalty of −11 and a gap-extension penalty of −1. A 9 × 15 position-pair scoring matrix was generated for each alignment, with rows corresponding to positions in the HLA-I 9mer and columns corresponding to positions in the HLA-II 15mer. For each pair of aligned amino acids, the corresponding BLOSUM62 substitution score was assigned to the respective matrix cell. Scores for cells corresponding to unaligned residue pairs or gaps were set to 0.

For each alignment method and HLA-II immunogenicity group, the position-pair scoring matrices were averaged element-wise across all selected epitope pairs. The Δscore matrix was calculated by subtracting the mean matrix for immunogenic HLA-I–non-immunogenic HLA-II pairs from that for immunogenic HLA-I–immunogenic HLA-II pairs (Eq. 11).

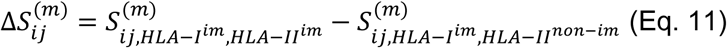

where i and j denote positions in the HLA-I 9-mer and HLA-II 15-mer, respectively, and (m) denotes the alignment method (local or global). The resulting 9 × 15 Δscore matrices were used to evaluate position-specific differences in sequence similarity.

Results from local alignment are presented in Figure 4B–E and Supplementary Figure 4E, whereas results from global alignment are presented in Supplementary Figure 4B–D and 4F–G. P-values were calculated using the two-sided Kolmogorov-Smirnov test.

### Integrated model from DON-B, DON-S, and DON-H2

DeepOmicsNeo-Immunogenicity (DON-IM) is designed to categorize immunogenic and non-immunogenic epitopes based on their binding affinity, and then apply fine-grained adjustments using the selfness and DON-H2. For the training and validation, we used a processed pHLA-I NCI immunogenicity dataset^16^ and IEDB-T^9^.

DON-IM receives as input the same N-dimensional feature set used by DON-B, augmented with the selfness score output from DON-S, yielding an (N+1)-dimensional input. Training was performed as a single-stage multi-task scheme rather than sequential distillation and fine-tuning stages: DON-IM was initialized from random weights and trained end-to-end to jointly learn pMHC-I binding affinity and immunogenicity. Two data sources contributed to this objective. Augmented samples drawn from the DON-B training set were labeled with DON-B’s own affinity predictions, providing a distillation-style auxiliary supervision signal for binding affinity, and were optimized using the conditional-MSE (cMSE) and weighted-binary-cross-entropy (wBCE) losses defined above. Independently, the immunogenicity training data - derived from the curated IEDB-T dataset^9^ together with an augmented set centered on IEDB-T-positive peptides - retained their native immunogenicity labels, which were used for the binary-cross-entropy-loss-with-logits (BCEWithLogitsLoss) and batch-group-wise-Bayesian-Personalized-Ranking (BPR) losses^29^, while additionally receiving DON-B-derived affinity pseudo-labels for the cMSE and wBCE losses. Consequently, DON-B-derived samples were supervised by cMSE and wBCE only, whereas IEDB-T-derived samples were jointly supervised by cMSE, wBCE, BCEWithLogitsLoss, and BPR to unify general binding-affinity learning with immunogenicity discrimination within a single multi-task training objective.

Because the alignment procedure employed by DON-H2 requires substantial CPU resources, incorporating DON-H2 as a conventional ensemble component was impractical. Instead, the ensemble score was computed by subtracting the DON-H2 output from the DON-IM output. The default configuration restricted DON-H2 inference to peptides from pHLA-I candidates with a DON-B percentile-rank below 2%.

Immunogenicity data curation and training strategy for DeepOmicsNeo-Immunogenicity Data augmentation was performed by generating hard-negative decoys from the human reference proteome, under the assumption that native human peptides are non-immunogenic under any condition. For each IEDB-T-positive peptide, candidate decoys were selected from the reference proteome within a defined maximum Hamming distance: up to 6 mismatched residues for 11mers and 12mers, 5 for 10mers, and 3 for 8mers and 9mers. Candidate decoys were then ranked by similarity to the corresponding positive peptide using a two-step procedure: peptides were first grouped by the number of matching residues (2, 1, or 0) at the canonical anchor positions (P2 and the C-terminus, PΩ), and within each group, further ordered by Hamming distance, such that the most similar decoys were prioritized as hard negatives. Each hard-negative peptide was assigned the same HLA allele as its corresponding positive peptide (e.g. a hard-negative derived from an HLA-A*02:01-restricted positive was likewise assigned HLA-A*02:01).

Because immunogenicity assays are performed under highly heterogeneous experimental conditions, positive and negative calls from different studies may not be directly comparable. To account for this, two preprocessing steps were applied prior to training. First, for deduplication, any pHLA-I allele combination with at least one positive report was labeled positive, irrespective of coexisting negative reports for the same combination. This step reflects the assumption that pHLA-I combinations reported as both positive and negative lie near the detection threshold, and would be expected to yield positive results under stronger immunogenic stimulation conditions. Second, studies sharing at least one common pHLA-I combination were grouped into connected experimental batches by transitive closure, using PubMed-IDentifiers (PMIDs) as study identifiers. Specifically, a graph was constructed in which each node represented a study (PMID) and an edge connected two studies if they shared at least one pHLA-I combination. Batch groups were then defined as the connected components of this graph. This grouping strategy was designed to restrict pairwise comparisons to studies linked through shared pHLA-I combinations, while avoiding the generation of an excessive number of redundant positive–negative pairs and reducing the computational complexity associated with pairwise ranking during model training. For example, if studies A (PMID001) and B (PMID002) shared at least one pHLA-I, and studies A and C (PMID003) also shared at least one pHLA-I, then A, B, and C were assigned to the same batch-group even if B and C did not directly share any pHLA-I. Conversely, studies with no shared immunogenic pHLA-I connections were assigned to separate batch groups. The primary determinant of peptide immunogenicity is consistently reported to be its binding affinity for the restricting HLA-I allotype. Motivated by this, and in contrast to prior approaches that frame epitope prediction purely as a binary classification problem, we jointly trained the model to predict binding strength and immunogenicity within a unified framework. Under this framework, we further assumed that among peptides reported as immunogenic, those with relatively stronger binding affinity constitute comparatively better epitopes.

Accordingly, the binding-dataset was optimized using cMSE and wBCE losses, while IEDB-T-derived immunogenicity samples were optimized using these same cMSE and wBCE losses together with an additional BCEWithLogitsLoss (Eq. 12) and, within each batch-group, a BPR loss (Eq. 13). The BPR loss was introduced as a supplementary term to further strengthen discrimination among samples that are more likely to be comparable, namely those originating from the same batch-group. Because training the model on the entire dataset at once is computationally impractical, the data are divided into small subsets called mini-batches, and the model’s parameters are updated after processing each mini-batch. Since a ranking loss compares samples against one another, it can only be computed among samples that appear together in the same mini-batch. The BPR loss was computed only for positive-negative pHLA-I pairs originating from the same batch-group and co-occurring within the same mini-batch. Specifically, for each batch-group represented in a mini-batch, all possible positive-negative pairs were constructed, and a pairwise ranking loss was computed using the softplus function applied to the temperature-scaled score difference between negative and positive samples, with a temperature parameter of T. The final training objective was defined as the sum of the BCEWithLogitsLoss loss and the BPR loss, where the BPR term was weighted by an epoch-dependent coefficient, (*λ*(t)). The coefficient followed a decay schedule, gradually reducing the influence of the BPR loss as training progressed.

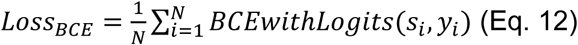

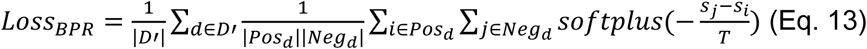

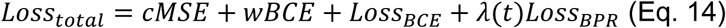

, where N denotes the number of samples in a mini-batch, *s_i_* and *y_i_* represent the predicted logit and binary label of the i-th sample, respectively, and D′ denotes the set of experimental dataset IDs for which both positive and negative samples are present within the same mini-batch. For each dataset d∈D′, *Pos_d_* and *Neg_d_* denote the sets of positive and negative samples, respectively. The BPR loss is computed by averaging the pairwise ranking loss over all positive-negative pairs within each dataset, where the pairwise score difference (*s_j_* −*s_i_*) is scaled by a temperature parameter T before applying the softplus function. Here, *s_i_* and *s_j_* correspond to the predicted logits of positive and negative samples, respectively. The mini-batch size was 16,384, and the BPR temperature (T) was set to 0.5. The BPR weight was initialized at 0.0625 and decreased according to a cosine schedule from epoch 0 to epoch 10, reaching a minimum weight of 0.

### Evaluation for TESLA, HiTIDE, and TiTANIAN benchmark

To evaluate model performance on experimentally validated data, we curated TESLA immunogenicity datasets from Tables S4 and S7, which we named TESLA-validated-benchmark-set (n = 911). To further evaluate model performance in a more clinically relevant setting, we generated a patient-derived neoantigen dataset from whole-exome sequencing (WES) data. Sequencing reads were aligned to the human reference genome (GRCh38) using BWA (v0.7.17)^30^, and somatic variants were identified with Mutect2 (v4.3.0)^31^. The coding effects of the identified variants were annotated using VEP (v101). HLA-I alleles were inferred using OptiType (v1.3.1)^32^, and candidate pHLA-Is were generated using the pVACseq pipeline (v2.0.4)^33^. For each epitope, immunogenicity scores were predicted against every HLA-I allele present in the corresponding patient’s genotype (typically up to six HLA-I alleles), and the highest score among these was assigned as the representative score for that epitope. This process yielded 41 immunogenic, 863 non-immunogenic, and 640,487 untested epitopes; untested epitopes were conservatively labeled as non-immunogenic. Model performance on this dataset was evaluated using AUC-ROC, PRAUC, and top-20 precision. The evaluation method is identically applied to the HiTIDE benchmark. In the case of TiTANIAN, we used TiTANIAN_IM_Neoantigen.csv^20^.

## Result

### Overall structure of DeepOmicsNeo models

We present three DeepOmicsNeo models for predicting the immunogenicity of peptides presented by MHC-I. In this workflow, DeepOmicsNeo-Binding (DON-B) predicts the binding affinity between candidate peptides and MHC-I (Figure 1A). DeepOmicsNeo-Selfness (DON-S) evaluates the degree to which candidate peptides resemble human peptides, which we will refer to as ‘selfness’. DeepOmicsNeo-Homology-with-HLA-II-epitopes (DON-H2) evaluates candidate peptides’ homology with known HLA-II-epitopes. Finally, DeepOmicsNeo-Immunogenicity (DON-IM) integrates these three models to estimate the immunogenic potential of candidate peptides. The following sections will detail the rationale behind our model architecture and performance.

### Pseudo-9mer profile is an effective representation for non-9mer peptide-MHC-I binding prediction

The first novel feature of our pMHC-I binding model DON-B is the embedding method for non-9mer peptides.

MHC-I molecules interact with 8 - 12 mer peptides, yet the vast majority of available binding data consists of 9mer peptides. Noting this, we hypothesized that providing 9mer interaction information when predicting non-9mer peptides would lead to a robust model for pMHC-I binding prediction. To this end, we introduced a method to transform non-9mer peptides into pseudo-9mer profiles using aligners. Our model takes the entire sequence of non-9mer peptides and the MHC-I pseudo-34mer through the aligner, and then generates bindable-9mers and non-bindable-9mers (Supplementary Figure 1A). Specifically, the aligner learns which parts of the peptide sequence need to be edited to maximize the Hadamard product of MHC-I’s PSSM for non-9mer peptides (see Methods for further detail). The positions to be edited are assigned probability values, which are used to create bindable- and non-bindable-pseudo-9mer profiles (Supplementary Figure 1A and 1B). This information is then passed to an affinity predictor to estimate binding affinity (Figure 1A).

To investigate whether the pseudo-9mers generated by the aligner actually produce meaningful bindable and non-bindable profiles, we predicted the binding affinities of pseudo-9mers edited from 1,000 non-9mer peptides for 15 different HLA-I alleles using NetMHCpan and MHCflurry, affinity predictors independent of our model. As expected, the bindable pseudo-9mers showed stronger binding affinity compared to the non-bindable pseudo-9mers in both programs (Figure 1B and Supplementary Figure 1C). Therefore, these results suggest that our pseudo-9mer serves as a suitable peptide representation for the non–9mer peptide-MHC-I binding prediction task.

For prediction of binding affinity for a non-9mer peptide, first the aligner generates non-bindable (nP9) and bindable pseudo-9mers (bP9) from the full peptide sequence (fP) and MHC-I sequence (M), then the affinity predictor predicts the pMHC-I affinity from them.

### pMHC-I-optimized data augmentation enhances pMHC-I binding prediction performance

The second novel feature of our binding model is the training data augmentation method. MHC-I interacts with peptides generally ranging from 8 to 12 mer. Despite this length variability, peptides binding to the same MHC-I allele often exhibit similar sequence motifs, particularly at anchor positions (Supplementary Figure 2A). We hypothesized that this characteristic could be leveraged for data augmentation of positive binder datasets. To validate this hypothesis, we first investigated whether peptides known to bind a specific MHC-I allele indeed display higher sequence similarity compared to random sequences, even when their lengths differ (Supplementary Figure 2B, see method). As anticipated, peptides known to bind the same MHC-I allele showed significantly higher sequence similarity to each other compared to random sequences, regardless of length differences (Figure 2B and Supplementary Figure 2C and 2D). Notably, when examining the frequency of edit and mismatch positions that yielded the maximum sequence alignment score against the most similar 9mer — or, for 9mer peptides themselves, against the most similar other 9mer within the same allele (excluding itself; see method) — we observed the lowest frequencies at the anchor positions, P2 and the C-terminus, for 8mer, 9mer, and 10mer peptides (Supplementary Figure 2E-G). This finding aligns with previous studies highlighting the critical role of anchor residues in peptide-MHC-I binding^34^. Based on these observations, we developed a novel data augmentation method for training K-mer prediction models.

**Figure S2.**
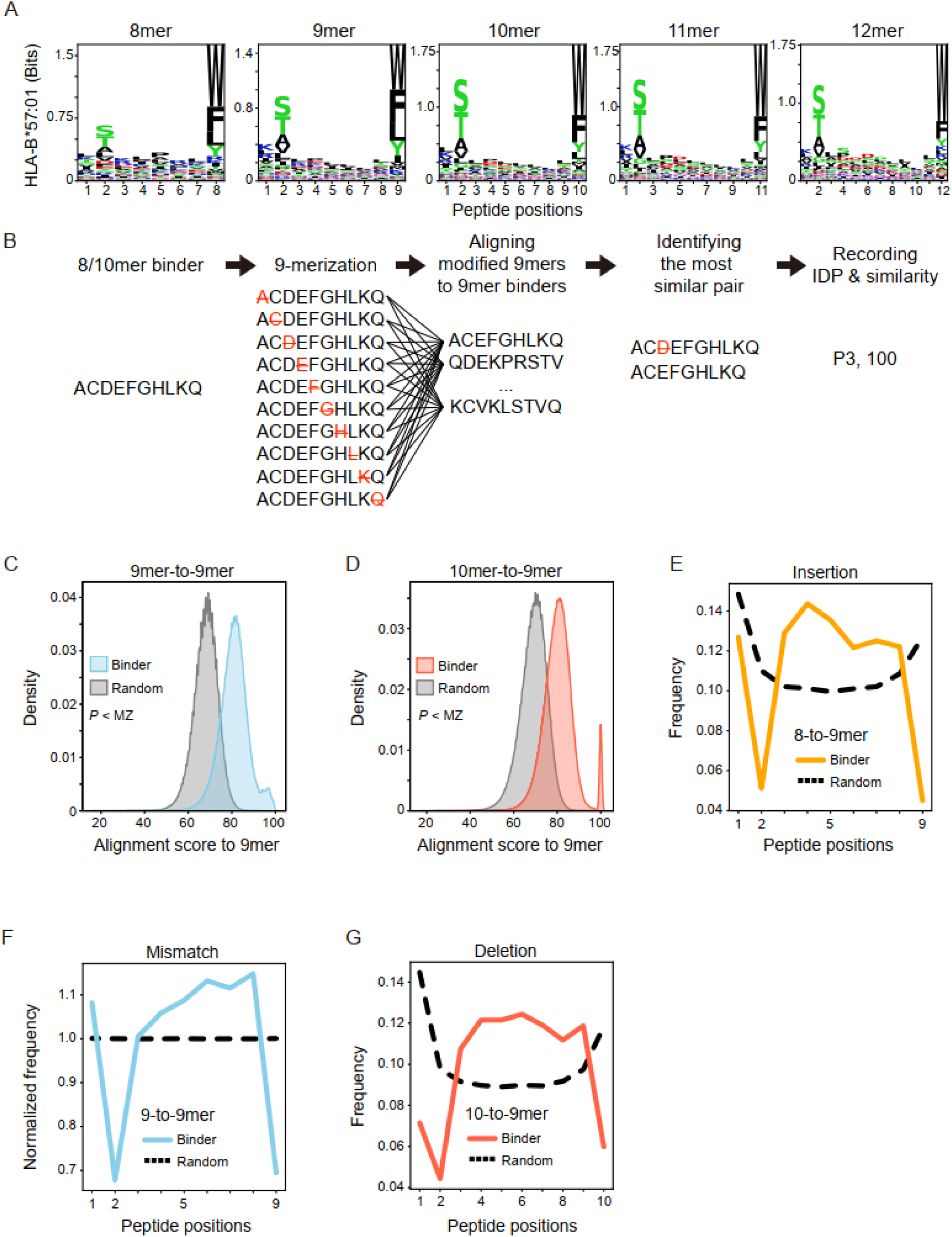
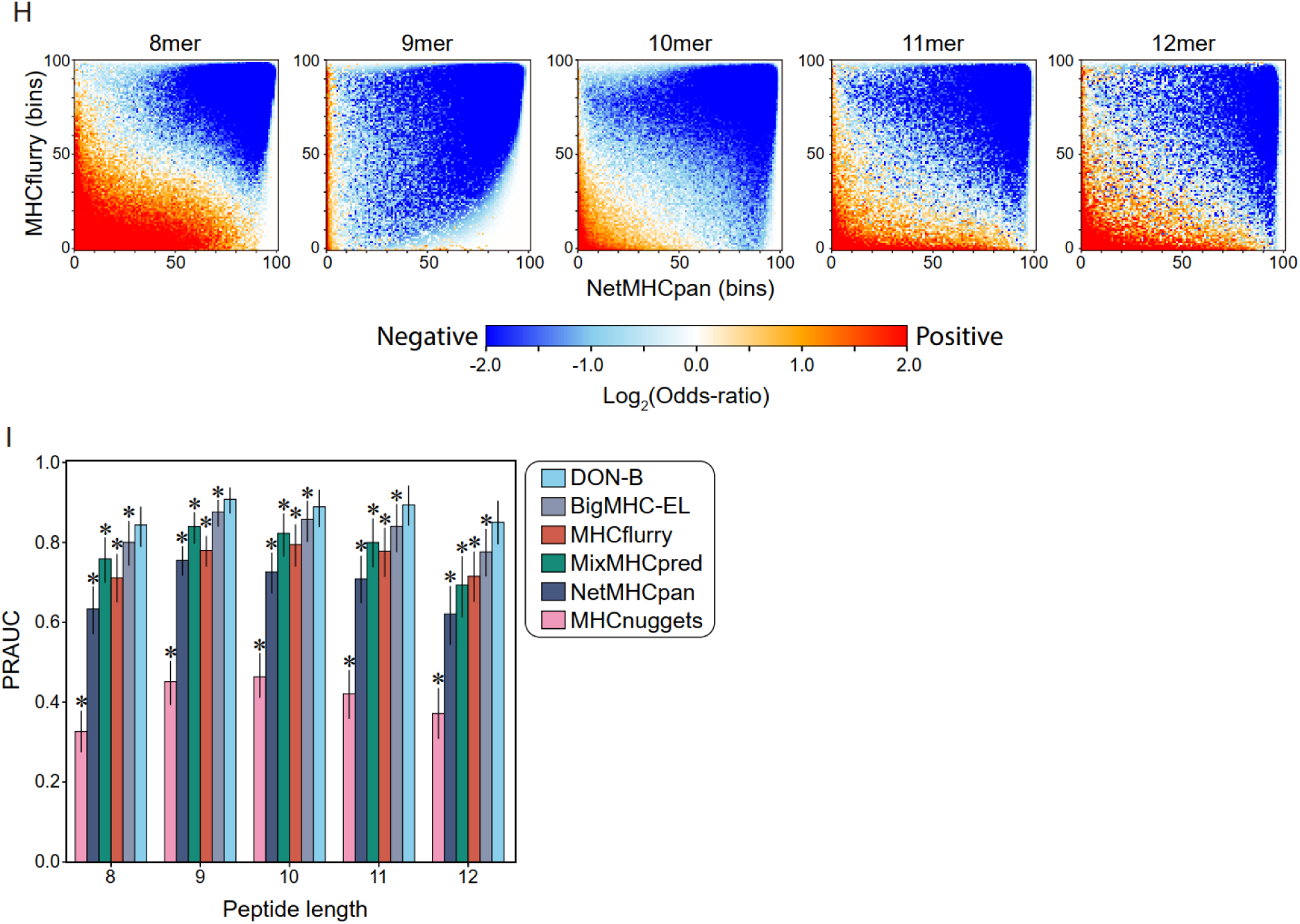
Details of DeepOmicsNeo-Binding and performance improvement in pMHC-I binding prediction task through data augmentation. A. Representative sequence logo of pMHC-I binding peptides. Different peptide lengths are indicated at the top. The X- and Y-axes represent peptide positions and Bits, respectively. Higher Bits scores indicate a higher preference for certain amino acids at the position. B. Workflow for comparing pMHC-I binding sequence similarity between different peptide lengths. Non-9mer pMHC-I binding peptide sequences are modified into 9mers by insertion or deletion. The modified peptides are aligned to actual pMHC-I binding 9mer peptides. Then, the highest alignment score and its modified position (IDP) are recorded. See Methods for details. C-D. Comparison of alignment scores between MHC-I-binding peptides and random peptides. For 9mer peptides (C), each binder’s alignment score was calculated against the most similar other 9mer sequence (excluding itself) known to bind the same MHC-I allele; for 10mer peptides (D), scores were calculated against the most similar 9mer sequence within the same allele. Score distributions of random peptides are shown in grey, while 9mer and 10mer binders are shown in blue and red, respectively. MZ represents machine-zero, indicating that the p-value was smaller than the smallest value representable in floating-point precision and therefore underflowed to zero. E-G. Frequency of insertion, mismatch, and deletion positions required to achieve the maximum alignment score against the most similar 9mer sequence binding the same MHC-I allele. The x-axis represents peptide positions, and the y-axis represents the frequency of modification at each position. The frequency distribution for random peptides is denoted by the dotted black line. In the case of mismatch (i.e., 9mer-to-9mer), frequency was normalized to the mean frequency across all positions within each group (binder or random) to account for their differing baseline scales. H. Distribution of predicted binding affinities of augmented pMHC-I pairs derived from the binders. Legends are as in Figure 2C. I. Comparison of six pMHC-I binding prediction algorithms in multi-allelic benchmark sets. Y-axis represents the Precision-Recall-AUC score. Legends are the same as in Figure 2E. P-values were obtained by the Wilcoxon signed-rank test. Exact p-values are provided in Supplementary Table 2.

Our strategy involves inserting, mutating and deleting amino acids from the P3 to P(ω-1) region of known positive binders, thereby generating synthetic training data while preserving the crucial anchor residues. Specifically, when building the K-mer affinity predictor, we first extracted only the MHC-I binding peptides from the K-1mer to K+1mer datasets. We then randomly modified the sequences at the inter-anchor positions. Using these modified sequences, we constructed a new dataset (referred to as the K’-mer dataset). To estimate plausible affinity of K’-mer sets, we trained Editor models with the real K-mer and random negative-sets (note: Editor model is essentially a DON-B model trained without the augmented data). The Editors were employed to label the affinity labels of the K’-mer dataset and negative-sets. Finally, the re-labeled datasets were combined with the original K-mer dataset and used to train the final K-mer model (Figure 2A, See methods). As independent validation of our approach, predicted binding strengths obtained with NetMHCpan and MHCflurry indicated that augmented pMHC-I dataset from the binder dataset exhibited significantly stronger affinities compared to random pMHC-I pairs (Figure 2C and Supplementary Figure 2H).

The model trained with the augmented data showed significantly better performance in 10-fold cross-validation compared to the model trained without augmentation (Figure 2D). The final DON-B model incorporating pseudo-9mer encoding method and trained with the augmented data demonstrated a superior performance (AUC: 0.962 ± 0.06 and PRAUC: 0.89 ± 0.14) over other state-of-the-art binding affinity prediction programs in the benchmark test dataset comprised of multi-allelic immunopeptidome data measured from 42 samples (Figure 2E, Supplementary Figure 2I, and Supplementary Table 2).

### GAN-based unsupervised anomaly detection can be used to predict immunogenicity

During the T-cell development process in the thymus, thymocytes remove a large portion of T-cells that have strong reactivity against self-peptides. The remaining T-cell repertoire is responsible for reacting to foreign non-self peptides, although they have never encountered or trained against them. Thus a model trained only with a human self-peptidome may be able to predict immunogenicity of a pMHC to a certain degree, in the context of self-tolerance.

Drawing parallels with the key concept of unsupervised anomaly detection, we employed a Generative Adversarial Network (GAN)-based anomaly detection method to mimic the negative selection in the thymus (Figure 3A). The GAN model consists of a generator and a discriminator. The generator’s role is to produce synthetic data that closely resembles self-human peptidome data, while the discriminator is tasked with distinguishing between genuine and fake data. Through an adversarial training process, these two models engage in a competitive learning dynamic. This adversarial interplay leads to the refinement of both models, with the generator continuously improving its ability to create realistic synthetic data, and the discriminator enhancing its capacity to accurately discern real from synthetic data.

**Figure 3.**
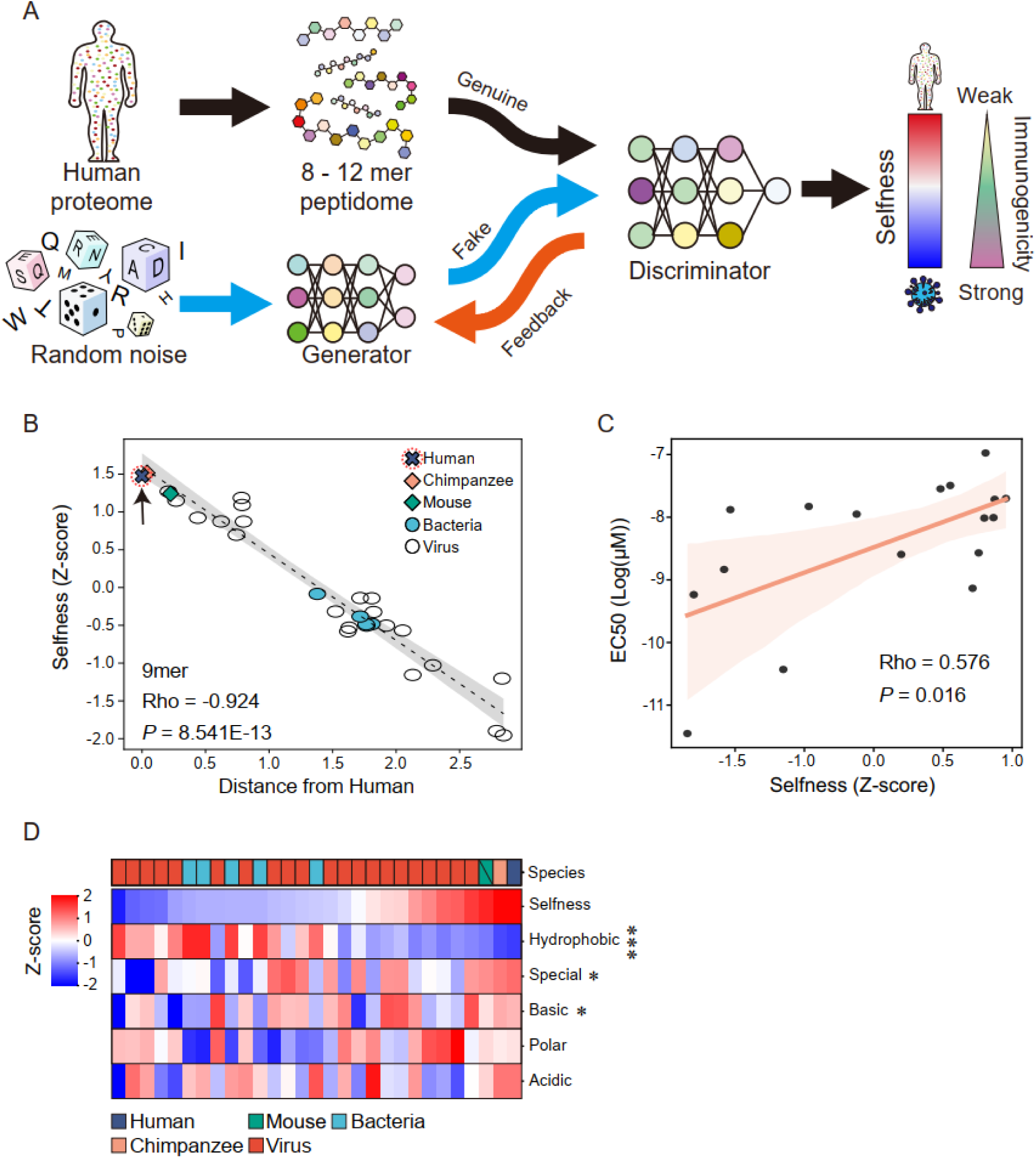
Development and characteristics of DeepOmicsNeo-Selfness. A. Overall workflow of the generation of non-immunogenic human self-peptidome (NIHSP) and training DeepOmicsNeo-Selfness. The number of human proteins and peptides are shown in parentheses. A higher selfness score is associated with a weaker immunogenic potential. B. The latent distances of 29 model organisms from Human (x-axis) and their selfness scores (y-axis). The colors and styles of points indicate the category of organisms. Human dot is highlighted by arrow and red-dotted circle. The statistical significance was evaluated by two-sided Spearman’s correlation test. C. The correlation between selfness score (x-axis) and T-cell functional avidity (y-axis). Their statistical significance was evaluated by two-sided Spearman’s correlation test. The avidity data was obtained from Schmidt et al.^22^. D. The correlation between selfness score and amino acid usage. Organisms are sorted according to their selfness scores. The statistical significance of correlation between amino acid group usage and selfness score was evaluated by two-sided Spearman’s correlation test. Asterisks indicate statistical significance (p < 0.05). Exact p-value is provided in Supplementary Table 3. (* : p<0.05, *** : p<0.001)

**Figure 4.**
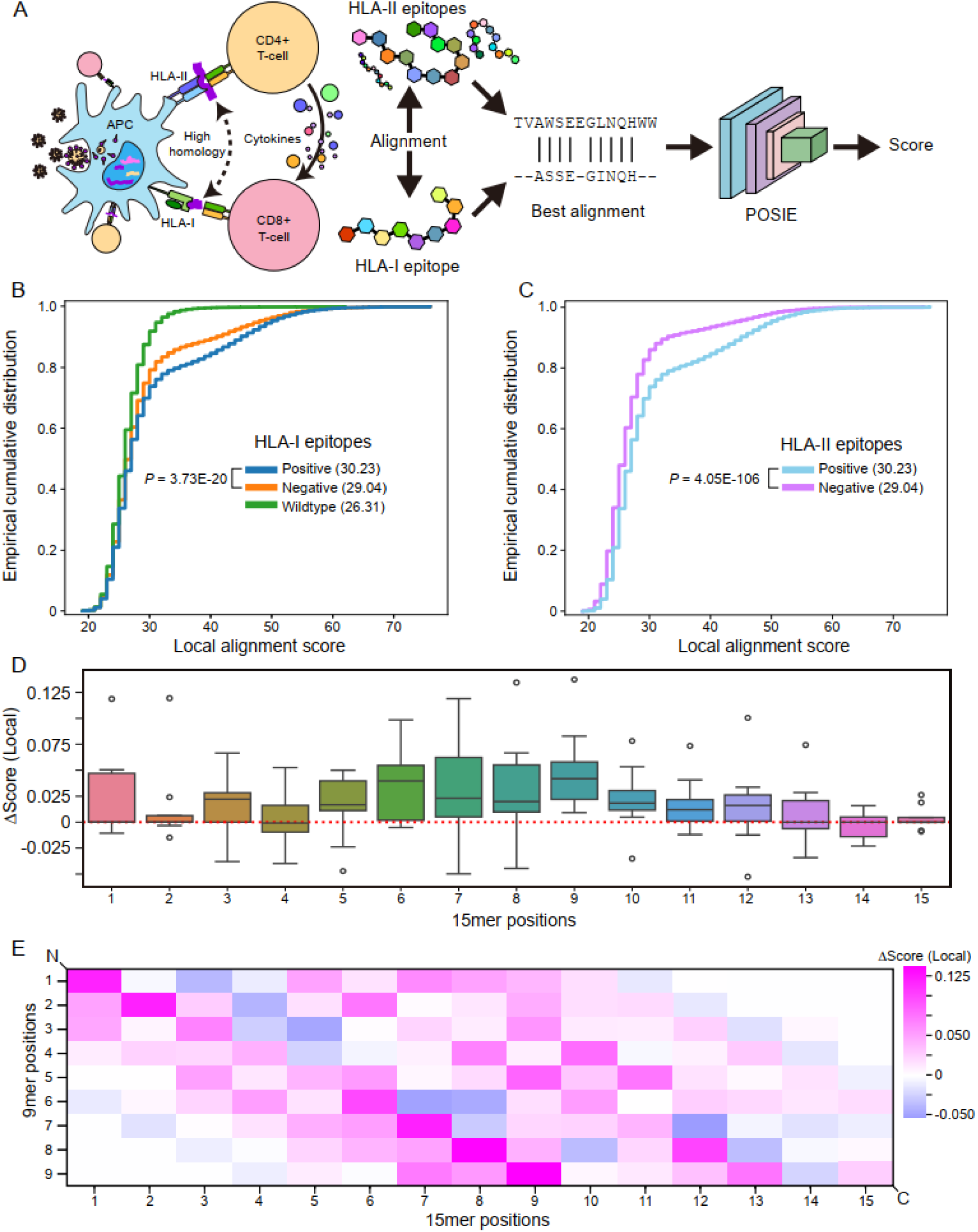
Patterns of shared immunogenic epitopes between the dual HLA classes. A. Overall workflow for DeepOmicsNeo-Homology-with-HLA-II-epitope (DON-H2). B. Distribution of local alignment scores between HLA-I epitopes and known immunogenic HLA-II epitopes. HLA-I epitopes are color-coded according to their immunogenicity status (immunogenic (blue), non-immunogenic (orange), and randomly selected human wildtype peptides (green)). Mean alignment scores are shown in parentheses. Statistical significance was evaluated by the two-sided Kolmogorov-Smirnov test. C. Distribution of local alignment scores between HLA-II epitopes and immunogenic HLA-I epitopes. HLA-II epitopes are color-coded according to their immunogenicity status. Mean alignment scores are shown in parentheses. P-values were calculated using two-sided Kolmogorov–Smirnov tests. D. The distribution of delta alignment scores per position in 15mer peptides. Higher Δscores indicate that the sequence alignment score for the amino acid at that position is higher in the immunogenic peptide compared to the non-immunogenic peptide. The dotted red line indicates 0. E. The joint distribution of delta alignment scores for 9 and 15mer peptides. N-terminus and C-terminus are marked by diagonal lines.

The ultimate outcome is a discriminator capable of recognizing the patterns of immunogenic peptides.

We created a human self-peptidome dataset by slicing 121,766 protein sequences into peptides ranging 8mer to 12mer, resulting in 57,683,512 peptide sequences (See method). Then, we trained the GAN models with the self-peptidome.

We first examined whether our GAN model (DeepOmicsNeo-Selfness, DON-S) could distinguish between self and non-self-peptidomes. We calculated the average peptidome profiles of 29 model organisms and their selfness scores (See method). As expected, the selfness scores for human and chimpanzee were the highest, while those for mouse, viruses and bacteria were significantly lower (Figure 3B and Supplementary Figure 3A). Additionally, a tendency was observed that as the latent distance from the human peptidome increased, the selfness score of the organism peptidome decreased (Figure 3B and Supplementary Figure 3A), indicating that our DON-S model can distinguish between self and non-self-peptidomes.

**Figure S3.**
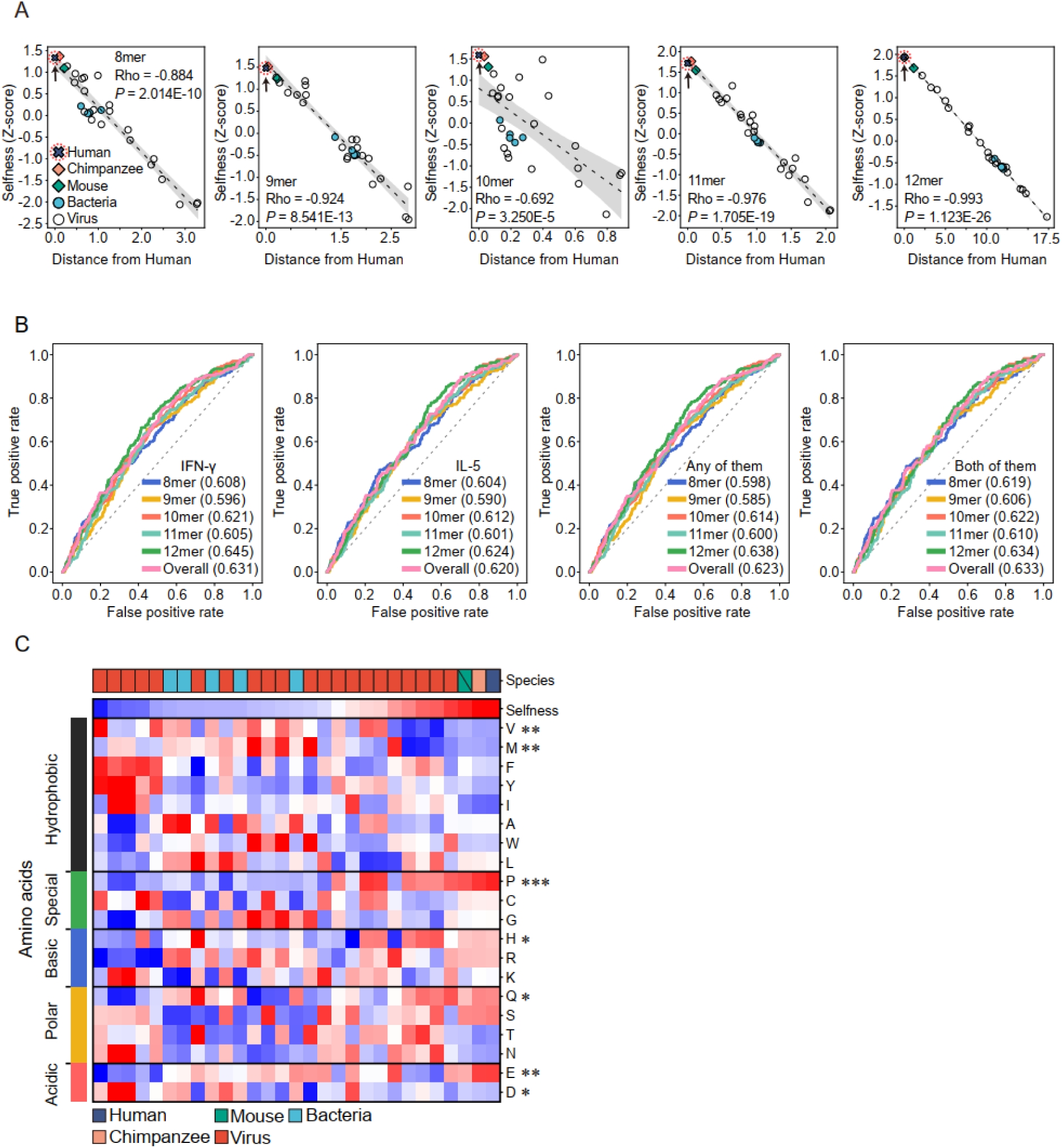
Characteristics of DeepOmicsNeo-Selfness. A. The latent distances of 29 model organisms from Human (x-axis) and their selfness scores (y-axis). The colors and styles of points indicate the category of organisms. The statistical significance was evaluated by two-sided Spearman’s correlation test. B. The performance of DeepOmicsNeo-Selfness in T-cell response prediction task. The names of the cytokines are displayed at the individual graphs. ‘Any of them’ refers to the prediction task of determining whether a peptide induces the secretion of either IFN-γ or IL-5. ‘Both of them’ refers to the prediction task of determining whether a peptide induces the secretion of both IFN-γ and IL-5 simultaneously. The colors of lines indicate the lengths of sliced peptides. Overall is obtained by average selfness scores of sliced peptides from the original peptides. The area under curve (AUC) values are shown in parentheses. C. The correlation between selfness score and amino acid usage. Organisms are sorted according to their selfness scores. The statistical significance of correlation between amino acid usage and selfness score was evaluated by two-sided Spearman’s correlation test. Asterisks indicate statistical significance (p < 0.05). Exact p-value is provided in Supplementary Table 3. (* : p<0.05, ** : p<0.01, *** : p<0.001)

**Supplementary Figure 4.**
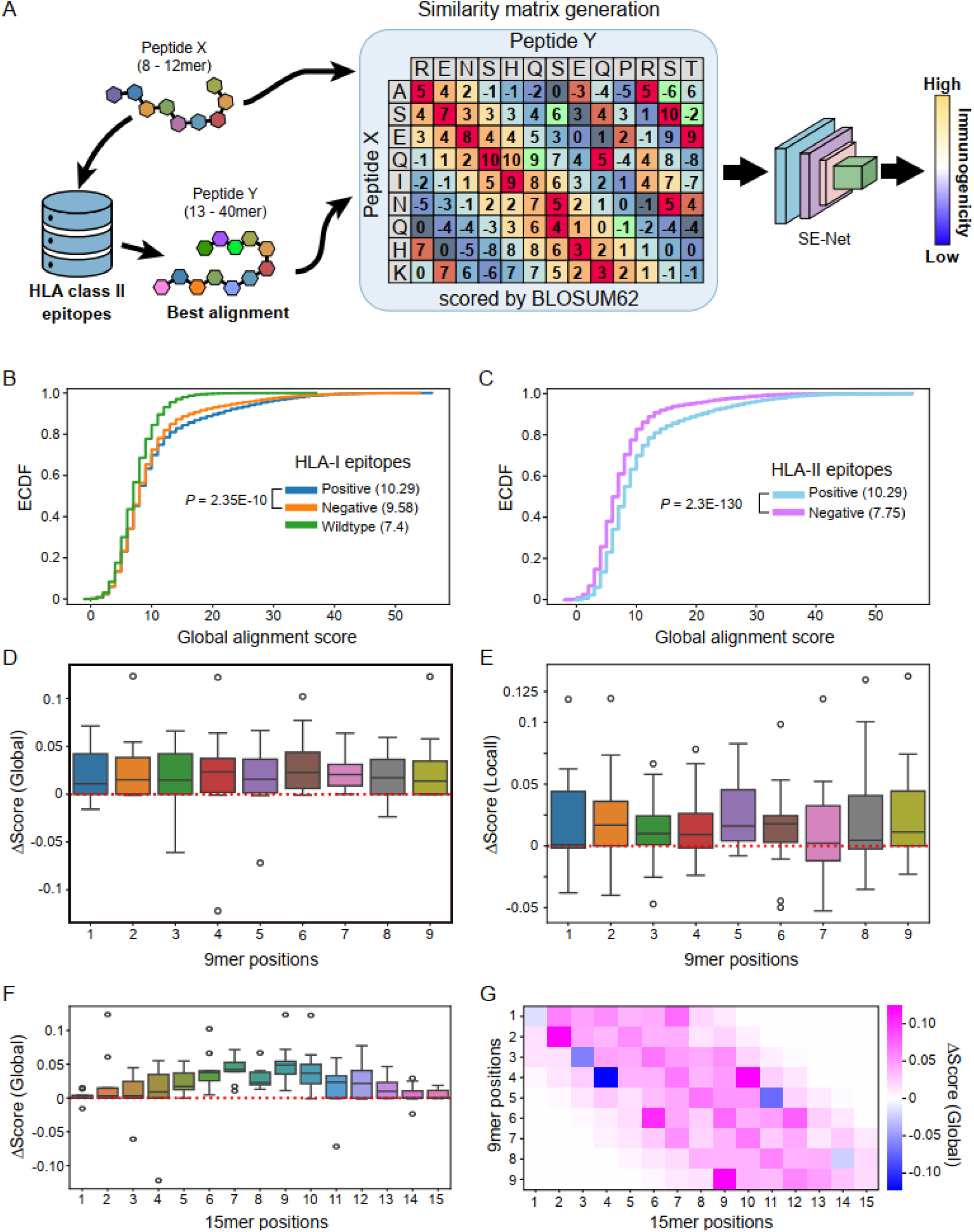
Patterns of shared immunogenic epitopes across the two HLA classes. A. Overview of the DeepOmicsNeo-Homology-with-HLA-II-epitope (DON-H2) architecture. For a given HLA-I epitope (Peptide X; 8–12-mer), the HLA-II epitope (Peptide Y; 13–40-mer) with the highest sequence-alignment score is retrieved from the HLA-II epitope database. An X-by-Y similarity matrix is then generated using BLOSUM62 substitution scores for each amino acid pair and provided as input to DON-H2, a squeeze-and-excitation network (SE-Net), which predicts an immunogenicity score. B. Distribution of global alignment scores between HLA-I epitopes and immunogenic HLA-II epitopes. HLA-I epitopes are color-coded according to their immunogenicity status. Mean alignment scores are shown in parentheses. C. Distribution of global alignment scores between HLA-II epitopes and immunogenic HLA-I epitopes. HLA-II epitopes are color-coded according to their immunogenicity status. Mean alignment scores are shown in parentheses. P-values were calculated using two-sided Kolmogorov–Smirnov tests. D–E. Distributions of positional relative alignment scores for immunogenic HLA-I 9-mer epitopes aligned to immunogenic and non-immunogenic HLA-II 15-mer epitopes. The y-axis indicates the position-specific relative alignment score, whereas the x-axis denotes the position within the HLA-I 9-mer epitope. The dotted red line indicates 0. The alignment method is indicated in parentheses on the y-axis. F. Position-specific relative global alignment scores of HLA-II epitopes. HLA-II epitopes were aligned to immunogenic HLA-I epitopes using global alignment. The legend is the same as that in Figure 4B. G. Position-specific relative global alignment scores between HLA-I and HLA-II epitopes. Sequences were aligned using global alignment. The legend is the same as that in Figure 4.

We next investigated whether the selfness score is also correlated with T-cell functional avidity and immunogenicity. The selfness score from DON-S was significantly correlated with T-cell functional avidity as measured by EC50 of IFN-γ ELISPOT assay reported by Schmidt et al. (Figure 3C). The selfness score was also correlated with T-cell immunogenicity as measured by ELISPOT assays for IFN-γ and/or IL-5 secretion reported by Miller et al. (Supplementary Figure 3B).

To decipher the scoring mechanism of DON-S, we investigated the correlation between amino acid usage per organism and the selfness scores. Regarding coefficients for amino acids, hydrophobic AAs which have been reported to be enriched in immunogenic epitopes^35,36^ generally exhibited strong negative values (Rho = -0.828, P-value = 3.068E-8), while other AAs showed weak positive values (Figure 3D and Supplementary Figure 3C).

Collectively, these results suggest that DeepOmicsNeo-Selfness recognizes some patterns of self-peptides, particularly hydrophobicity, which are related to T-cell mediated immune response.

### High homology with immunogenic epitopes of HLA-II provides informative clues for predicting HLA-I immunogenicity

Studies have demonstrated the necessity of help from CD4+ T-cells in activation of CD8+ T-cells against tumor neoantigens, and effective cancer vaccines are those in which the MHC-I and MHC-II epitopes reside on the same peptide strand, potentially leading to more efficient uptake and presentation of both antigens by the same antigen-presenting cells^37,38^ (Figure 4A). Halpert et al. further reported that the co-expression of MHC-I and II epitopes with overlapping sequences in dendritic cells triggers PAMP-like response to strongly induce Th1 immune polarization^39^.

Therefore, we hypothesized that HLA-I epitopes with high similarity to known HLA-II epitopes would be more immunogenic than the ones without correspondence.

To validate our hypothesis, we compared the sequence similarity between HLA-I-immunogenic-peptides and HLA-II-immunogenic-peptides against HLA-I-nonimmunogenic-peptides and random 8 - 12mer sequences from the human wildtype proteome. Strikingly, HLA-I-immunogenic-peptides exhibited significantly higher sequence similarity to HLA-II-immunogenic-peptides compared to non-immunogenic sequences (Figure 4B and Supplementary Figure 4B). Furthermore, HLA-I-immunogenic peptides showed higher sequence similarity to HLA-II-immunogenic peptides than to HLA-II-non-immunogenic peptides (Figure 4C and Supplementary Figure 4C).

Next, we examined sequence alignment patterns using 9mer and 15mer peptides, which represent the most abundant peptide lengths for HLA-I and HLA-II, respectively. For 9mer immunogenic peptides, 15mer immunogenic peptides displayed higher alignment scores than 15mer non-immunogenic peptides regardless of position (Supplementary Figure 4D and E). However, analysis of the distribution of alignment scores within 15mer peptides revealed that while similarity at the N-to-N or C-to-C terminal ends was low, similarity increased toward the central region (Figure 4D-E and Supplementary Figure 4F-G). These results suggest that HLA-I-epitopes are more likely to be immunogenic when they share high sequence similarity with the central portion of immunogenic HLA-II-epitopes.

Based on these findings, we developed DeepOmicsNeo-Homology-with-HLA-II-epitope (DON-H2) that leverages these patterns of epitopes between dual-HLA classes. Given a query HLA-I-epitope, the DON-H2 model identifies homologous HLA-II-immunogenic-peptides and predicts the immunogenicity of HLA-I-epitope based on BLOSUM62 amino-acid substitution score matrix of two sequences (Figure 4A and Supplementary Figure 4A, See method).

### Development of the immunogenicity prediction model (DON-IM) integrating DON-B, DON-S, and DON-H2

In CD8+ T-cell immune responses, the stable binding of pHLA-I complexes, the non-selfness of peptides, and high homology with HLA-II may all contribute. Therefore, to develop an accurate neoantigen prediction model, we constructed an integrated model by combining DON-B, DON-S, and DON-H2 (Figure 5A, See Methods).

**Figure 5.**
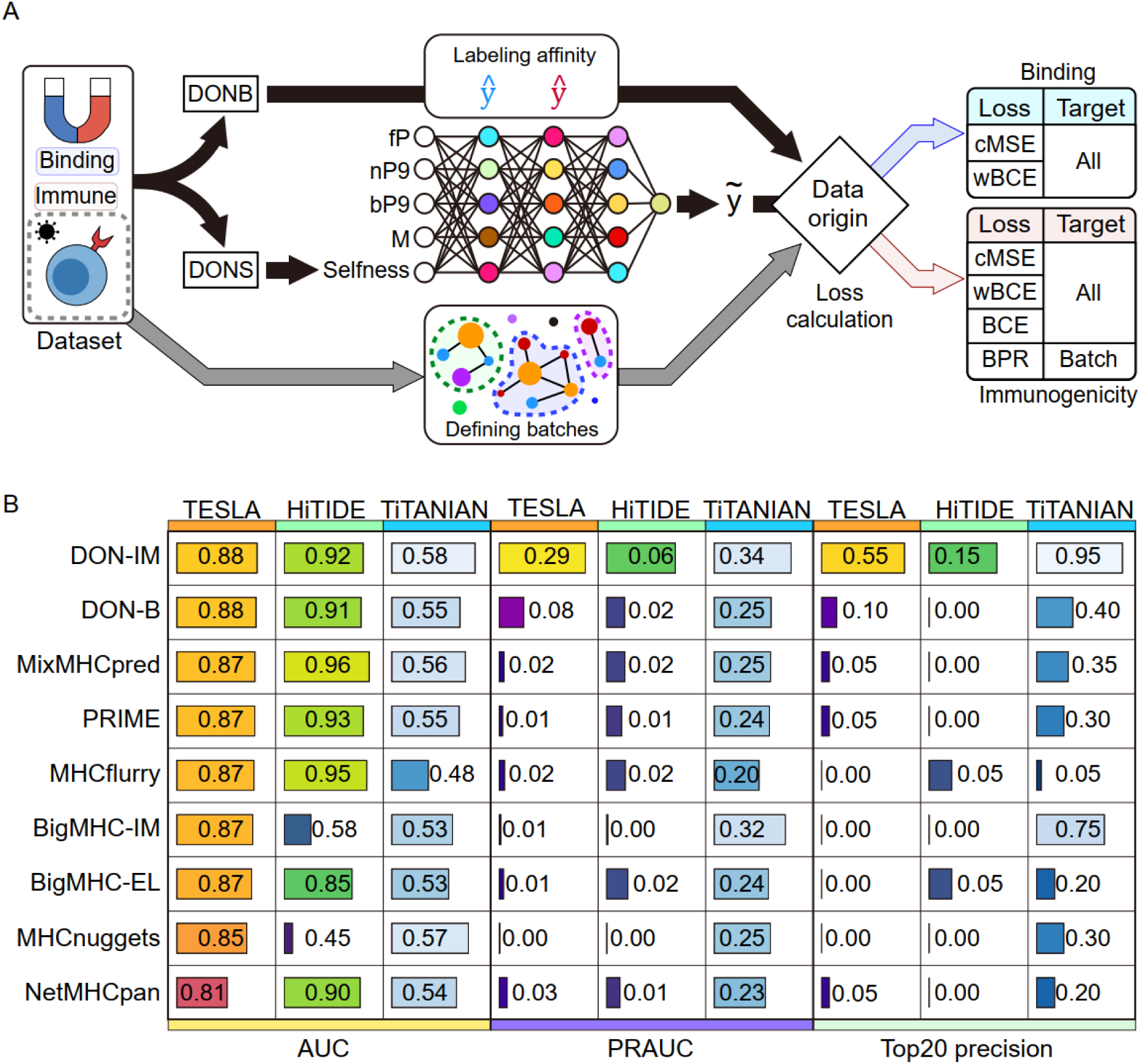
Development and performance of DeepOmicsNeo-Immunogenicity. A. Schematic training workflow of DeepOmicsNeo-Immunogenicity (DON-IM). B. Performance evaluation of DON-IM and DON-B with seven published predictors on the TESLA, HiTIDE, and TiTANIAN benchmark sets. Predictors are listed by row. Each benchmark set is shown as a column block, and each block reports three metrics: AUC (area under the receiver operating characteristic curve), PRAUC (area under the precision-recall curve), and Top20 precision (fraction of true positives among the 20 highest-scoring peptides). Bars and numerical values indicate the metric value for each predictor. Cell shading indicates relative performance: color intensity is scaled separately for each combination of benchmark set and metric, and brighter shades denote higher values.

Immunogenicity labels are essentially binary, yet they are compiled from assays performed under highly heterogeneous experimental conditions, so positive and negative calls from different studies are not directly comparable. We therefore assumed that, even among peptides reported as immunogenic, epitopes with stronger binding affinity are of higher quality than those with weaker binding affinity. Based on this assumption, DON-IM was trained end-to-end within a single multi-task framework that jointly learned binding affinity and immunogenicity, rather than through sequential pretraining and fine-tuning stages.

Affinity-related supervision was derived from the same training data employed for DON-B, relabeled with continuous binding-affinity scores predicted by DON-B as soft targets, and optimized using the same loss functions applied to DON-B. For immunogenicity data, we redesigned both the data curation procedure and the training method. For data curation, hard-negative decoys were sampled from human wild-type reference sequences, with selection biased toward peptides sharing the same residues as the corresponding positive peptide at the P2 and C-terminal anchor positions (see Methods for further details).

In addition, because batch effects across studies may compromise model generalization, we added a batch-group-wise Bayesian-personalized-ranking (BPR) loss to the standard binary-cross-entropy (BCE) loss. Datasets were first partitioned by PubMed-IDentifier (PMID), and studies sharing at least one pHLA-I combination were grouped into the same batch-group via transitive closure. During training, all data points within a mini-batch contributed to the BCE loss, whereas the BPR loss was applied exclusively between pairs originating from the same batch-group. This design was intended to sharpen the model’s discrimination between immunogenic and non-immunogenic epitopes within each batch-group (Supplementary Figure 5A-C; see Methods).

**Supplementary Figure 5.**
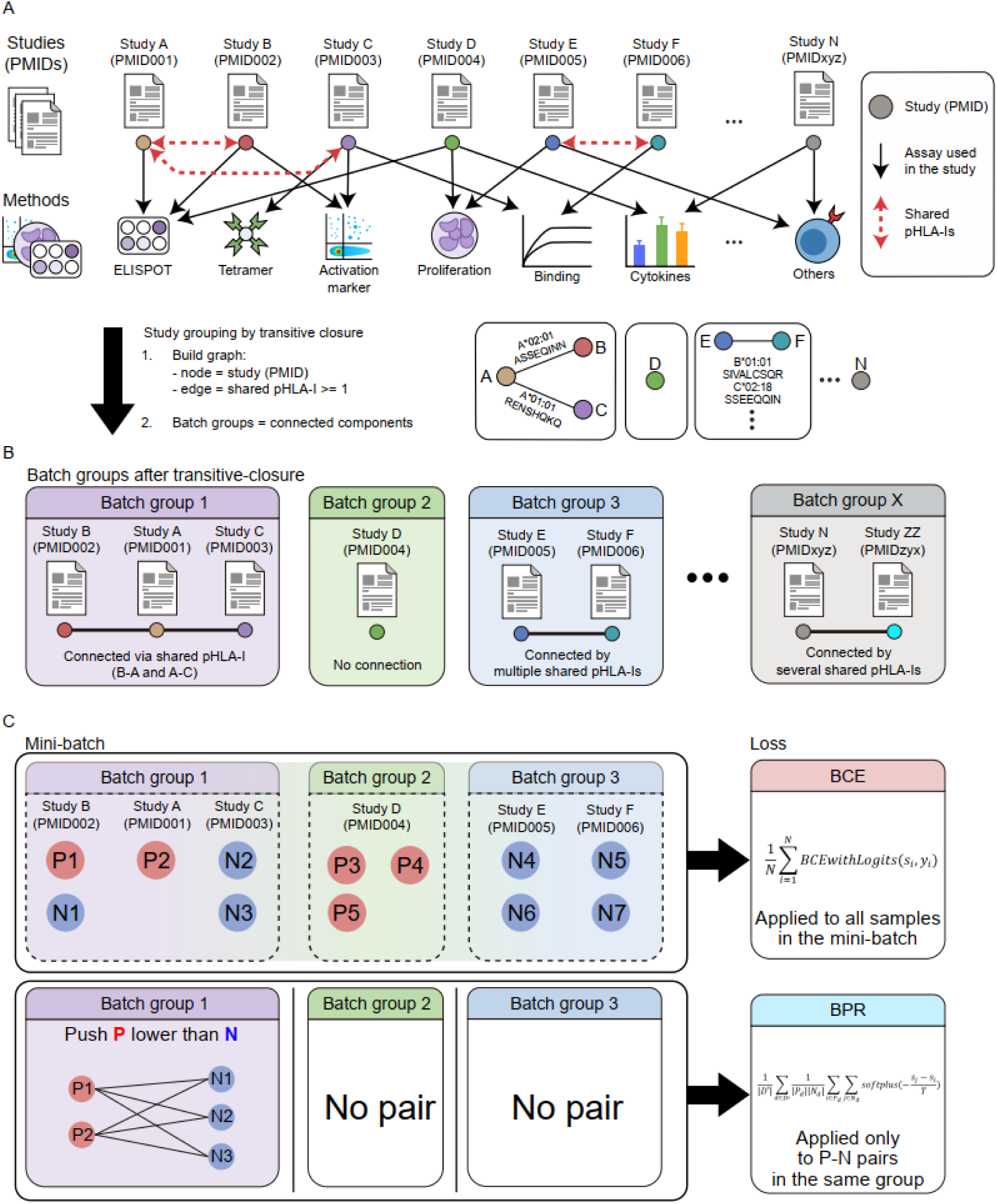
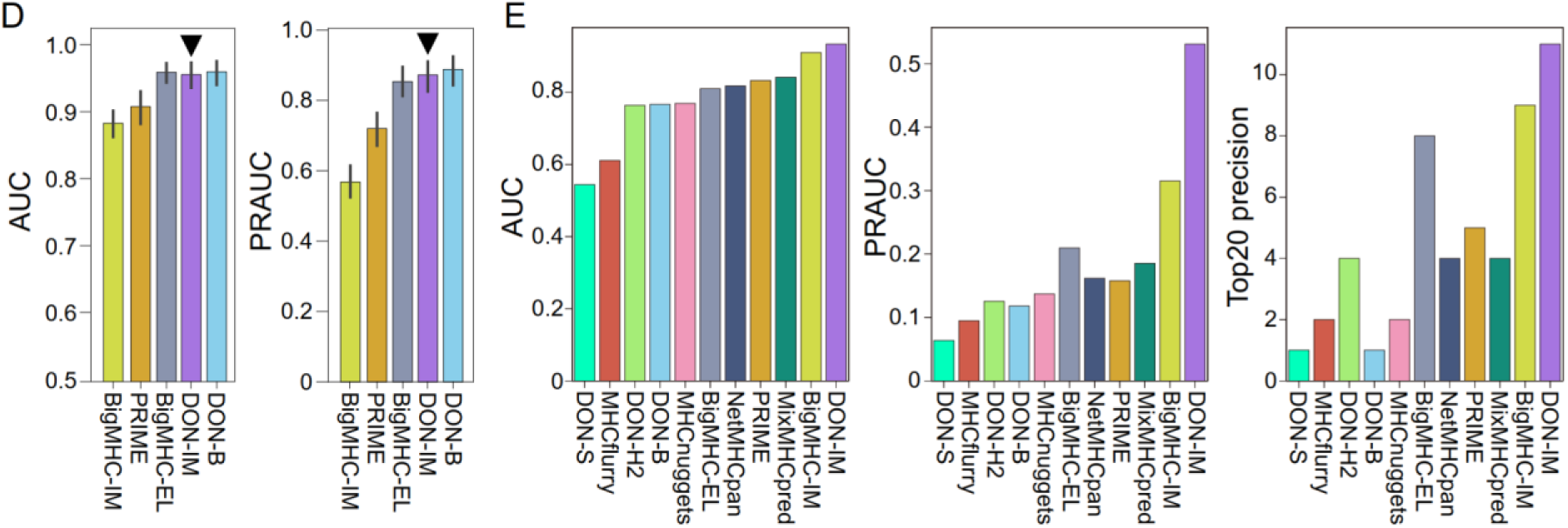
Batch-clustering and training strategy for DeepOmicsNeo-Immunogenicity. A. Heterogeneity and connectivity of the training data. The assay used in each study is represented by black arrows, and pHLA-I epitopes shared between studies are represented by dotted red arrows. A graph is constructed in which each node represents a study and each edge represents one or more shared pHLA-I epitopes between two studies (see example subgraphs). B. Clustering of studies into batch groups. Studies sharing one or more pHLA-I epitopes are grouped into a batch group via transitive-closure. Studies without any shared epitope form a singleton batch group (e.g. Study D). C. Application of loss functions during training. (Upper) All samples within a mini-batch are used to compute the binary-cross-entropy (BCE) loss, regardless of batch group. (Lower) Only samples from batch groups containing both positive and negative pairs within a mini-batch are used to compute the Bayesian-personalized-ranking (BPR) loss. D. Peptide and MHC class I binding prediction performance of DON-IM. DON-IM is highlighted by reverse-triangles. Legend is the same as Figure 2E. E. Performance evaluation of DeepOmicsNeo models with seven published predictors on the TESLA-validated-benchmark-set (n = 911). AUC, PRAUC, and Top20 precision are shown on the y-axis.

DON-IM showed a strong performance on pHLA-I binding prediction compared to the other immunogenicity and binding prediction models (AUC: 0.958 ± 0.06 and PRAUC: 0.872 ± 0.144) (Supplementary Figure 5D). In addition DON-IM demonstrated improved predictive performance compared to the parental models in TESLA-validated-benchmark-set (Supplementary Figure 5E). For personalized cancer vaccine design, proportion of true positives among selected top 20 candidate neoantigens (top20 precision) is the most clinically relevant benchmark when comparing neoantigen prediction algorithms. When cases from TESLA benchmark dataset were analyzed starting from sequencing data, top20 precision of DON-IM was 55% (11/20 immunogenic) compared to only 5% (1/20 immunogenic) for the next best performing algorithms (MixMHCpred, PRIME, and NetMHCpan) (Figure 5B). Although not ideal, previous studies using TESLA benchmark started their analyses with TESLA preselected and experimentally tested dataset (N=911), which tends to inflate the result due to preselection bias. DON-IM outperformed SOTA models even in such an analysis setting (Top20 precision of 55% vs 45%, PRAUC 0.55 vs 0.3) (Supplementary Figure 5E). For HiTIDE dataset, Top20 precision for DON-IM and the best performing competitors (MHCflurry and BigMHC-EL) were 15% and 5% respectively. For TiTANIAN dataset, Top20 precision for DON-IM was 95% compared to 75% for BigMHC-IM.

Collectively, these results suggest that integrating pMHC-I binding and immunogenicity enables DON-IM to robustly prioritize immunogenic neoantigens across diverse benchmarks.

## Discussion

In this paper, we introduced DeepOmicsNeo, which is inspired by biological principles in the T-cell immune system.

For DeepOmicsNeo-Binding, we devised a novel peptide encoding method, pseudo-9mer, to deliver interaction characteristics of peptide-MHC-I (pMHC-I) effectively when training non-9mer models. Our study shows that the proposed methods indeed can improve the performance of pMHC-I binding prediction.

Data augmentation is an important technique in machine learning that enhances the size and diversity of training datasets by generating synthetic or modified data samples from existing ones. This approach is essential for improving model generalization and performance, particularly in scenarios with limited or imbalanced data^40^.

For nearly two decades, the training of pMHC-I binding prediction models has relied on augmenting negative binding datasets by combining random peptides with MHC-I molecules, generating datasets 20 - 100 times larger than experimentally validated data^23,41–45^. This approach has effectively reduced potential biases in training data while providing diverse pMHC-I data points, thereby enhancing model performance. Paradoxically, however, the selective and high-affinity interactions between MHC-I molecules and a small subset of peptides have posed significant challenges for developing effective augmentation strategies for positive binders, which are more critical for identifying immunogenic peptides in applications such as neoantigen-based cancer immunotherapy where precision is more important than specificity.

To address this challenge, we developed a novel positive binder augmentation method that introduces random sequence variations in non-anchor positions while preserving the critical anchor positions. This approach statistically significantly improved model performance compared to the model trained without positive binder augmentation. Our method complements the established random pMHC-I negative binder augmentation by enriching the diversity of positive binders, thus improving the model’s ability to distinguish strong binders from non- and weak-binders.

Despite its success, our method has a limitation. The reliance on predefined anchor positions may overlook allele-specific nuances in sub-binding motifs^46^, potentially limiting generalizability to less-studied MHC-I alleles. Therefore, future work should explore more efficient ways to consider sub-anchor positions.

We developed DeepOmicsNeo-Selfness by leveraging the similarities between anomaly detection and the negative selection of T-cells. To our knowledge, our report is the first to demonstrate the application of unsupervised anomaly detection machine-learning techniques in the immunogenicity prediction task. Our results show that DeepOmicsNeo-Selfness reflects important patterns in immunogenic epitopes. Notably, the correlation between the selfness score and the proportion of amino acids aligns with reports from not only molecular experiments^35,36^ but also other neoantigen prediction models grounded in supervised learning techniques^24,47,48^. This alignment suggests a potential solution to address the data scarcity issue in T-cell reactivity data, especially regarding true negatives^49^.

HLA-I immunogenicity has traditionally been modeled using pHLA-I binding affinity and intrinsic epitope features such as hydrophobicity and proteasomal cleavage^50^. However, CD8+ T-cell activation also depends on coordinated interactions among antigen-presenting cells, which use HLA-II systems. Earlier attempts to incorporate peptide-HLA-II interactions did not establish a predictive link to HLA-I immunogenicity, possibly because the two classes differ markedly in binding propensities. We show that immunogenic HLA-I epitopes share significantly higher sequence homology with immunogenic HLA-II epitopes than with non-immunogenic peptides, consistent with evidence that MHC-I epitope immunogenicity is influenced by homologous immunogenic epitopes in the MHC-II repertoire^39^. Notably, MHC-I epitopes aligned preferentially with the central region of MHC-II epitopes rather than the N- or C-terminus. This is consistent with the structure of the MHC-II groove, which is open-ended and anchors peptides mainly through a central 9mer core, leaving the termini largely non-interacting^51,52^. The central enrichment in 15mers may therefore reflect constraints of MHC-II binding stability and concurrent presentation of overlapping epitopes during T-cell priming.

By formalizing these cross-class relationships, DON-H2 shows that shared immunogenic patterns across the two HLA classes help identify immunogenic HLA-I epitopes. However, alignment is computationally costly, so DON-H2 must be applied to a pre-filtered set of high-affinity binders; more efficient algorithms are needed to exploit this shared immunogenicity. To accurately recapitulate the complexity of CD8+ T-cell immune responses, we constructed DeepOmicsNeo-Immunogenicity (DON-IM), a framework that integrates three potential determinants of immunogenicity: pMHC-I binding, self-tolerance, and epitope homology with HLA-II. Rather than combining independently trained models post hoc, DON-IM incorporates binding- and self-tolerance-related information directly within a single end-to-end network, through multi-task supervision distilled from DON-B and architectural integration of DON-S, and further refines this output with a DON-H2-derived homology signal to yield the final immunogenicity score (see Methods). By harmonizing these complementary immunological signals, DON-IM demonstrated superior predictive performance over its individual parental components on the diverse benchmarks.

Beyond this architectural integration, we also addressed a data-level challenge specific to immunogenicity modeling. Prior computational approaches have typically relied on standard binary-cross-entropy (BCE) loss applied to immunogenicity labels drawn from a single experimental assay type (e.g. ELISPOT), implicitly treating positive and negative calls as directly comparable regardless of their study of origin. However, immunogenicity datasets compiled across heterogeneous experimental conditions and cohorts can introduce substantial batch effects that confound genuine biological signal with assay- or study-specific artifacts. To address this, we assumed that, among peptides reported as immunogenic, epitopes with stronger binding affinity are of higher quality, and we trained DON-IM to learn binding affinity jointly with immunogenicity, so that the model receives a graded signal in addition to the binary labels. In addition, DON-IM was trained with a batch-group-wise Bayesian-personalized-ranking (BPR) loss alongside BCE: studies sharing at least one common pHLA-I combination were grouped via transitive-closure into batch-groups, and ranking supervision was restricted to peptide pairs originating from the same batch-group. This design was intended to further sharpen discrimination between immunogenic and non-immunogenic epitopes within experimentally comparable contexts, rather than forcing comparisons across potentially incompatible assay conditions.

DON-IM provided the best top20 precision in all three tested benchmark datasets, significantly outperforming competing models, suggesting its clinical value in designing personalized cancer vaccines. It is worthwhile to note that in many published studies TESLA benchmark data were analyzed starting from the tested candidate epitopes rather than from the sequencing data. Since these tested epitopes were originally preselected by mostly MHC affinity based prediction programs, the analysis starting from tested epitopes may bias the results in a way to inflate the precision. We performed our testing starting from the sequencing data to avoid such bias. However, since untested candidate epitopes were regarded as negative, there might be false negatives in our analysis deflating our estimate for its performance.

In conclusion, these findings underscore that integrating binding, selfness, and helper-epitope homology provides a more reliable computational representation of pHLA-I immunogenicity and offers a robust framework for prioritizing neoantigens in precision immunotherapy. By formalizing this integration into a unified prediction framework, DeepOmicsNeo offers a better path toward identifying therapeutically actionable neoantigens in personalized cancer immunotherapy.

## Competing interests

Sejin Oh, Hae-Suk Kim, and Soonmyung Paik are employees of Theragen Bio Co., Ltd. The authors declare that they have no other competing interest.

## Author’s Contributions

Sejin Oh conceptualized the study, developed models, conducted the data analysis, and wrote the manuscript. Seong-Eui Hong and Hae-Suk Kim were involved in data curation. Soonmyung Paik supervised the study design, data analysis, and manuscript preparation. All authors read and approved the final manuscript.

## Acknowledgments

This research was supported by a grant of Korean ARPA-H Project through the Korea Health Industry Development Institute (KHIDI), funded by the Ministry of Health & Welfare, Republic of Korea (grant number : RS-2025-25402727).

## References

1. Weber, J. S. et al. Individualised neoantigen therapy mRNA-4157 (V940) plus pembrolizumab versus pembrolizumab monotherapy in resected melanoma (KEYNOTE-942): a randomised, phase 2b study. The Lancet 403, 632–644 (2024).

2. Lopez, J. et al. Autogene cevumeran with or without atezolizumab in advanced solid tumors: a phase 1 trial. Nat. Med. 31, 152–164 (2025).

3. Wells, D. K. et al. Key Parameters of Tumor Epitope Immunogenicity Revealed Through a Consortium Approach Improve Neoantigen Prediction. Cell 183, 818–834.e13 (2020).

4. Zyrin, V. et al. Evaluation of Computational Neoantigen Prioritization Strategies for Personalized Cancer Vaccines. Vaccines (Basel). 14, 857 (2026).

5. Jobin, K. et al. A distinct priming phase regulates CD8 T cell immunity by orchestrating paracrine IL-2 signals. Science 388, eadq1405 (2025).

6. Yan, Z. et al. Next-generation IEDB tools: A platform for epitope prediction and analysis. Nucleic Acids Res. 52, W526–W532 (2024).

7. Spindler, M. J. et al. Massively parallel interrogation and mining of natively paired human TCRαβ repertoires. Nat. Biotechnol. 38, 609–619 (2020).

8. McMaster, B., Thorpe, C., Ogg, G., Deane, C. M. & Koohy, H. Can AlphaFold’s breakthrough in protein structure help decode the fundamental principles of adaptive cellular immunity? Nat. Methods 21, 766–776 (2024).

9. Vita, R. et al. The Immune Epitope Database (IEDB): 2024 update. Nucleic Acids Res. 53, D436–D443 (2025).

10. Abelin, J. G. et al. Mass Spectrometry Profiling of HLA-Associated Peptidomes in Mono-allelic Cells Enables More Accurate Epitope Prediction. Immunity 46, 315–326 (2017).

11. Faridi, P. et al. A subset of HLA-I peptides are not genomically templated: Evidence for cis- and trans-spliced peptide ligands. Sci. Immunol. 3, 1–12 (2018).

12. Sarkizova, S. et al. A large peptidome dataset improves HLA class I epitope prediction across most of the human population. Nat. Biotechnol. 38, 199–209 (2020).

13. Abelin, J. G. et al. Defining HLA-II Ligand Processing and Binding Rules with Mass Spectrometry Enhances Cancer Epitope Prediction. Immunity 51, 766–779.e17 (2019).

14. Pyke, R. M. et al. Precision neoantigen discovery using large-scale immunopeptidomes and composite modeling of MHC peptide presentation. Molecular and Cellular Proteomics 20, (2021).

15. Yates, A. D. et al. Ensembl 2026. Nucleic Acids Res. 54, D1053–D1060 (2026).

16. Müller, M. et al. Machine learning methods and harmonized datasets improve immunogenic neoantigen prediction. Immunity 56, 2650–2663.e6 (2023).

17. Kumar, P. et al. Tumour-associated antigenic peptides are present in the HLA class I ligandome of cancer cell line derived extracellular vesicles. Immunology 166, 249–264 (2022).

18. Qi, Y. A. et al. Proteogenomic analysis unveils the HLA class I-presented immunopeptidome in Melanoma and EGFR-Mutant Lung Adenocarcinoma. Molecular and Cellular Proteomics 20, 100136 (2021).

19. Bassani-Sternberg, M. et al. Direct identification of clinically relevant neoepitopes presented on native human melanoma tissue by mass spectrometry. Nature Communications 7, (2016).

20. Kim, J. et al. T-SCAPE: T cell immunogenicity scoring via cross-domain aided predictive engine. Science Advances 11, 1–22 (2025).

21. Miller, A. M. et al. A functional identification platform reveals frequent, spontaneous neoantigen-specific T cell responses in patients with cancer. Sci. Transl. Med. 16, eabj9905 (2024).

22. Schmidt, J. et al. Neoantigen-specific CD8 T cells with high structural avidity preferentially reside in and eliminate tumors. Nat. Commun. 14, 3188 (2023).

23. O’Donnell, T. J. et al. MHCflurry: Open-Source Class I MHC Binding Affinity Prediction. Cell Syst. 7, 129–132.e4 (2018).

24. Schmidt, J. et al. Prediction of neo-epitope immunogenicity reveals TCR recognition determinants and provides insight into immunoediting. Cell Rep. Med. 2, 100194 (2021).

25. Huang, G., Sun, Y., Liu, Z., Sedra, D. & Weinberger, K. Q. Deep networks with stochastic depth. Lecture Notes in Computer Science (including subseries Lecture Notes in Artificial Intelligence and Lecture Notes in Bioinformatics) 9908 **LNCS**, 646–661 (2016).

26. Park, H. J. et al. HLA allele frequencies in 5802 Koreans: Varied allele types associated with SJS/TEN according to culprit drugs. Yonsei Med. J. 57, 118–126 (2016).

27. UniProt Consortium. UniProt: the Universal Protein Knowledgebase in 2023. Nucleic Acids Res. 51, D523–D531 (2023).

28. Hu, J., Shen, L., Albanie, S., Sun, G. & Wu, E. Squeeze-and-Excitation Networks. IEEE Trans. Pattern Anal. Mach. Intell. 42, 2011–2023 (2020).

29. Rendle, S., Freudenthaler, C., Gantner, Z. & Schmidt-Thieme, L. BPR: Bayesian personalized ranking from implicit feedback. Proceedings of the 25th Conference on Uncertainty in Artificial Intelligence, UAI 2009 452–461 (2009).

30. Li, H. & Durbin, R. Fast and accurate short read alignment with Burrows-Wheeler transform. Bioinformatics 25, 1754–1760 (2009).

31. Benjamin, D. et al. Calling Somatic SNVs and Indels with Mutect2. bioRxiv 861054 (2019).

32. Szolek, A. et al. OptiType: Precision HLA typing from next-generation sequencing data. Bioinformatics 30, 3310–3316 (2014).

33. Hundal, J. et al. pVAC-Seq : A genome-guided in silico approach to identifying tumor neoantigens. Genome Med. 1–11 (2016) doi:10.1186/s13073-016-0264-5.

34. Capietto, A. H. et al. Mutation position is an important determinant for predicting cancer neoantigens. Journal of Experimental Medicine 217, (2020).

35. Chowell, D. et al. TCR contact residue hydrophobicity is a hallmark of immunogenic CD8+ T cell epitopes. Proc. Natl. Acad. Sci. U. S. A. 112, E1754–E1762 (2015).

36. Calis, J. J. A. et al. Properties of MHC Class I Presented Peptides That Enhance Immunogenicity. PLoS Comput. Biol. 9, (2013).

37. Bennett, S. R. M., Carbone, F. R., Karamalis, F., Miller, J. F. A. P. & Heath, W. R. Induction of a CD8+ cytotoxic T lymphocyte response by cross-priming requires cognate CD4+ T cell help. Journal of Experimental Medicine 186, 65–70 (1997).

38. Alspach, E. et al. MHC-II neoantigens shape tumour immunity and response to immunotherapy. Nature 574, 696–701 (2019).

39. Halpert, M. M. et al. MHC class I and II peptide homology regulates the cellular immune response. FASEB J. 34, 8082–8101 (2020).

40. Mumuni, A. & Mumuni, F. Data augmentation: A comprehensive survey of modern approaches. Array 16, (2022).

41. Nielsen, M. & Andreatta, M. NetMHCpan-3.0; improved prediction of binding to MHC class I molecules integrating information from multiple receptor and peptide length datasets. Genome Med. 8, (2016).

42. Nielsen, M. et al. NetMHCpan, a method for quantitative predictions of peptide binding to any HLA-A and -B locus protein of known sequence. PLoS One 2, (2007).

43. Dönnes, P. & Elofsson, A. Prediction of MHC class I binding peptides, using SVMHC. BMC Bioinformatics 3, 1–8 (2002).

44. Albert, B. A. et al. Deep neural networks predict class I major histocompatibility complex epitope presentation and transfer learn neoepitope immunogenicity. *Nat*. Mach. Intell. 5, 861–872 (2023).

45. Shao, X. M. et al. High-Throughput Prediction of MHC Class I and II Neoantigens with MHCnuggets. Cancer Immunol. Res. 8, 396–408 (2020).

46. Jackson, K. R. et al. Charge-based interactions through peptide position 4 drive diversity of antigen presentation by human leukocyte antigen class I molecules. PNAS Nexus 1, 1–16 (2022).

47. Bravi, B. et al. A transfer-learning approach to predict antigen immunogenicity and T-cell receptor specificity. Elife 12, (2023).

48. Zhou, C. et al. pTuneos: prioritizing tumor neoantigens from next-generation sequencing data. Genome Med. 11, 67 (2019).

49. Buckley, P. R. et al. Evaluating performance of existing computational models in predicting CD8+ T cell pathogenic epitopes and cancer neoantigens. Brief. Bioinform. 23, (2022).

50. Kim, S. et al. Neopepsee: Accurate genome-level prediction of neoantigens by harnessing sequence and amino acid immunogenicity information. Annals of Oncology 29, 1030–1036 (2018).

51. Sarri, C. A. et al. Amino acid signatures in the HLA class II peptide-binding region associated with protection/susceptibility to the severe West Nile Virus disease. PLoS One 13, 1–20 (2018).

52. Liu, X. et al. Alternate interactions define the binding of peptides to the MHC molecule IAb. Proc. Natl. Acad. Sci. U. S. A. 99, 8820–8825 (2002).

